# The BELL-type homeobox transcription factor RLC3/OsBLH4 controls leaf rolling and drought tolerance via KNOX-BELL-lignin regulatory network in rice

**DOI:** 10.64898/2026.02.03.701930

**Authors:** Lei Qiao, Zhongyu Zhang, Qinying Li, Kairui Deng, Jinrong Li, Mingjia Lin, YuGuo Chen, Zhen Li, Lihong Zheng, Haifeng Li, Kunming Chen, Wenqiang Li

**Affiliations:** State Key Laboratory for Crop Stress Resistance and High-Efficiency Production, College of Life Sciences, Northwest A&F University, Yangling 712100, Shaanxi, China; College of Life Sciences, Xinyang Normal University, Xinyang 464000, Henan, China

**Author notes:** Author for correspondence: *Wenqiang Li,. These authors contributed equally to this work (Lei Qiao, Zhongyu Zhang).

**Keywords:** BELL-KNOX complex, drought tolerance, leaf rolling, lignin biosynthesis, rice (*Oryza sativa* L.), RLC3/OsBLH4

## Abstract

- Moderate leaf rolling in rice is crucial for plant architecture and stress adaptation, but its molecular regulation remains unclear. We investigated the role of RLC3/OsBLH4, a BELL-type homeobox transcription factor, in controlling leaf rolling and drought tolerance, addressing gaps in lignin biosynthesis and cell wall development mechanisms.
- We used gene map-based cloning (*rlc3-1*, *rlc3-2*), CRISPR/Cas9 knockout lines (*rlc3-ko#11*, *rlc3-ko#12*), and allelic complementation to validate *RLC3*’s function. Additionally, we employed biochemical assays, gene expression analysis, and protein interaction studies to explore its regulatory network.
- *RLC3* mutations impaired lignin biosynthesis and secondary cell wall formation, reducing bulliform cells area and causing midrib defects. These structural abnormalities accelerated water loss, leading to excessive leaf rolling and compromised drought tolerance. Mechanistically, RLC3 directly activates lignin synthesis genes (*OsPAL5*, *OsCOMT5*, OsCCR4, *OsCAld5H1*) and interacts with KNOX transcription factors (OSH1, OSH45, OSH71) to form a KNOX-BELL complex, further regulating lignin content and cell wall development.
- RLC3 orchestrates lignin deposition and secondary cell wall development to control leaf rolling, water transport, and drought tolerance. This study reveals a novel KNOX-BELL-lignin regulatory module governing leaf morphology and stress adaptation, offering targets for crop improvement under drought conditions.

## Introduction

Leaves critically determine plant architecture, photosynthetic efficiency, and stress resilience, directly influencing rice yield and grain quality (Lang *et al*., 2004a; You *et al*., 2022; Xu *et al*., 2023). Leaf morphology is mainly defined by three key parameters: leaf length, width, and rolling degree. Moderate rolling enhances leaf erectness and compactness, improving light transmittance and energy utilization (Lang *et al*., 2004a,b; Shen *et al*., 2009). It also reduces transpiration and thermal absorption, boosting stress tolerance (Kadioglu and Terzi, 2007; Zhang *et al*., 2018; Wang *et al*., 2023). Collectively, these physiological benefits elucidate the prevalence of moderate leaf rolling in modern rice varieties, particularly in high-yielding hybrid cultivars (Yuan, 1997, 2018; Shen *et al*., 2009; Chen *et al*., 2010). In contrast, excessive leaf rolling invariably triggers growth suppression, developmental aberrations, and substantial yield penalties, underscoring the critical importance of balanced morphological regulation in rice production systems.

Due to its agronomic importance in rice, leaf rolling has been extensively studied, with over 30 regulatory genes identified (Xu *et al*., 2018; Wang *et al*., 2020; Jia *et al*., 2024). This trait arises from multifactorial mechanisms involving epidermal cell patterning dysregulation, polarity disruption, aberrant differentiation, cell wall modifications, and physiological homeostasis fluctuations (Xu *et al*., 2018; Wang *et al*., 2020). Notably, most leaf-rolling phenotypes correlate with developmental abnormalities in bulliform cells (BCs). The rice *ROC5* gene, encoding a type IV HD-ZIP transcription factor, regulates BCs development: its loss-of-function mutation increases BC size and number, inducing abaxial rolling, while overexpression reduces these parameters, causing adaxial rolling (Zou *et al*., 2011). Similarly, genes such as *OsZHD1* (Xu *et al*., 2014), *ADL1* (Hibara *et al*., 2009), *ACL1* and *ACL2* (Li *et al*., 2010), *LC2* (Zhao *et al*., 2010), *REL2* (Yang *et al*., 2016), *LRRK1* (Zhou *et al*., 2018), *OsHox32*/*OSHB4* (Li *et al*., 2016; Zhang *et al*., 2018) and *OsSNF7.2* (Zhou *et al*., 2023) modulate leaf rolling by altering BCs morphology, proliferation, and/or spatial arrangement. Additionally, cell wall modifications critically influence leaf rolling. *OsOFP2-KNAT7/BLH6* suppresses lignin, inducing rolling via vascular defects (Schmitz *et al*., 2015), while *SRL1* and *RENL1* co-regulate cellulose biosynthesis, affecting rolling and drought tolerance (Liu *et al*., 2024). *RL14* modulates leaf rolling by altering secondary cell wall formation (Fang *et al*., 2012). Our studies show *CLD1/SRL1* loss causes bulliform-like epidermal cell expansion (increased water loss) and cellulose/lignin imbalance, driving leaf rolling (Li *et al*., 2017). These findings indicate that cell wall composition alterations disrupt secondary wall formation and hydraulic homeostasis, collectively driving BCs morphological changes and leaf rolling. Beyond its role in leaf rolling regulation, the cell wall’s structural composition, plays a pivotal role in drought resistance and overall stress adaptation (Malacarne *et al*., 2024). Specifically, lignin deposition in secondary cell walls not only strengthen structural integrity against water deficit but also activates antioxidant systems to scavenge reactive oxygen species (ROS) generated during drought stress, thereby synergistically enhancing plant survival under arid conditions (Bang *et al*., 2019, 2022; Hou *et al*., 2022). In maize, *ZmCAD* and *ZmCOMT* regulate lignin content via enzymatic activity (Hu *et al*., 2009). In rice, transcription factor OsNAC5 activates *OsCCR10*, a rate-limiting enzyme in monolignol biosynthesis, thereby promoting lignin accumulation in roots and enhancing drought tolerance (Bang *et al*., 2022). Additionally, transcription factors such as OsTF1L (Bang *et al*., 2019), OsERF71 (Lee *et al*., 2016) and OsNAC17 (Jung *et al*., 2022) orchestrate drought responses by binding to lignin biosynthesis genes or modulating their expression, thereby fine-tuning cell wall fortification under water deficit conditions. These studies demonstrate that the structural composition of cell walls, particularly lignin biosynthesis, critically contributes to both plant tissue development and morphogenesis, while also significantly enhancing drought stress resistance.

In higher plants, KNOTTED1-like homeobox (KNOX) proteins are critical transcriptional regulators that maintain meristem cell identity; however, their ectopic or prolonged expression disrupts the balance between meristem maintenance and organ differentiation, resulting in sustained SAM activity and aberrant leaf morphogenesis (Hay and Tsiantis, 2009). As TALE-family members, KNOX and BELL proteins share conserved domains and form heterodimers essential for developmental regulation (Bellaoui *et al*., 2001; Smith and Hake, 2003; Mukherjee *et al*., 2009). Systematic screens reveal promiscuous KNOX–BELL interactions in *Arabidopsis*, where all 13 BELLs bind KNOXs to sustain meristem function (Kumar *et al*., 2007); in potato, POTH1–StBEL5 represses *GA20ox1* (Chen *et al*., 2004); in soybean, GmBLH4–GmSBH1 mediates stress responses (Tao *et al*., 2018); and in maize, BLH12/14–KN1 controls inflorescence architecture (Tsuda *et al*., 2017). In rice, OSH15–qSH1/SH5 suppresses *CAD2* to reduce lignin and promote grain shattering (Yoon *et al*., 2017), while qSH1–OSH71 activates *OsXTH12* to modulate cell wall remodeling, thereby promoting grain shattering (Wu *et al*., 2025). Collectively, these findings highlight the evolutionarily conserved role of BELL–KNOX heterodimers in coordinating meristem activity, organ morphogenesis, and stress adaptation across monocots and dicots.

Although BELL-KNOX complexes regulate downstream targets and influence diverse developmental processes, the precise molecular mechanisms by which they coordinately control leaf morphology and stress adaptation remain incompletely understood. Here, we identify RLC3/OsBLH4, a BELL-type homeobox transcription factor that interacts with KNOX proteins (OSH1, OSH45, OSH71), as a key regulator of rice leaf rolling and drought tolerance. RLC3 directly targets lignin biosynthesis genes (*OsPAL5*, *OsCOMT5*, *OsCCR4* and *OsCAld5H1*), thereby modulating secondary cell wall deposition and water transport, which jointly determine leaf morphology and drought resilience. Our findings reveal a novel KNOX-BELL-lignin regulatory network, where RLC3 acts as a key regulator linking leaf development and stress adaptation, making it a prime target for rice improvement.

## Material and Methods

### Plant materials and growth conditions

The *rlc3-1* and *rlc3-2* mutants were isolated from an ethyl methanesulfonate (EMS)-induced mutant population of rice Nipponbare (Nip, *Oryza sativa* L. ssp. *japonica*). CRISPR/Cas9-mediated genome editing (Ma and Liu, 2016) was employed to generate knockout lines (*rlc3-ko#11, rlc3-ko#25*, *osh1-ko*, *osh45-ko*, and *osh71-ko*) in wild-type Nip and Zhonghua11 (ZH11, *Oryza sativa* L. ssp. *japonica*), respectively. Transgenic plants were produced via *Agrobacterium*-mediated transformation of rice calli (Toki *et al*., 2006). Field cultivation was conducted in experimental field at Mianxian, Shaanxi Province of China, and in pots at Northwest A&F University (Yangling, Shaanxi, China).

### Map-based cloning and allelic test

For genetic mapping, the *rlc3-1* and *rlc3-2* mutants were crossed with Kasalath (*Oryza sativa* L. ssp. *indica* cultivar), respectively. F₂ individuals exhibiting leaf rolling phenotypes were selected for DNA extraction and genetic mapping. Initial screening employed 254 SSR markers distributed across all 12 chromosomes, followed by fine-mapping using newly developed InDel markers. To identify the mutation site, genomic DNA fragments of the candidate gene were amplified from Nipponbare (Nip), *rlc3-1*, and *rlc3-2*. For sequence verification, all amplicons were Sanger-sequenced and aligned to the Nipponbare reference genome (IRGSP-1.0). Allelic tests were performed by pairwise crosses among *rlc3-1*, *rlc3-2*, and *rlc3-ko#25* mutants. Hybrid seeds from these crosses were sown to evaluate F₁ phenotypes, and DNA was extracted from hybrid plants for genotyping to confirm allelic relationships.

### Leaf traits analysis

For assessing leaf-related physiological traits, leaves from tillering-stage plants or heading-stage plants were used to determine the leaf rolling index (LRI) using the method described by Shi *et al*. (2007). To evaluate water-related stress responses, the second fully expanded leaves from rice plants at tillering stage were employed for measuring relative water content (RWC) and water loss rate (WLR) following the protocol described by Zou *et al*. (2011) and Mao *et al*. (2012), respectively.

### Drought stress assays

Drought stress experiments were conducted at seedling and reproductive stages. For the seedling-stage assay, plants were grown under were grown in a controlled environment (16-h light (30°C)/8-h dark (26°C) photoperiod, 180 µmol m⁻² s⁻¹ photosynthetic photon flux density, and 70–80% relative humidity) for 25 days before drought stress was imposed by withholding water for 3–7 days until visible wilting occurred. Subsequently, re-watering was performed for 4–5 days, and survival rates were calculated based on the proportion of plants that recovered after rehydration. For the reproductive-stage assay, an equal number of healthy seedlings from wild type and mutants were transplanted into soil pots and cultivated under natural field conditions until reaching reproductive maturity. Drought stress was then applied by withholding water for 3 days until wilting symptoms appeared, followed by 7 days of re-watering. Survival rates were recorded as described above. Physiological stress indicators such as chlorophyll content, malondialdehyde (MDA) levels, proline accumulation, and electrolyte leakage were measured in leaves from both control and drought-treated plants using the method described by Yang *et al*. (2006).

### Microscopic and transmission electron microscopic (TEM) observations

The second fully expanded leaves from tillering-stage plants were collected and immediately fixed in FAA solution (70% ethanol, 5% formaldehyde, and 5% acetic acid) at 4°C overnight for paraffin section preparation. Cross-sections were microtomed, stained, and analyzed using an Olympus BX51 microscope (Tokyo, Japan) equipped with a digital imaging system. For TEM analysis, fresh transverse sections of the second leaves were subjected to vacuum infiltration and fixed in a solution containing 2% (v/v) paraformaldehyde and 2.5% (v/v) glutaraldehyde overnight at 4°C. Sample preparation followed the protocol outlined by Li *et al*. (2009). Micrographs were acquired using a transmission electron microscope (TECNAI G2 SPIRIT BIO, FEI).

### Histological and histochemical analyses

To evaluate leaf structural integrity, toluidine blue (TB) staining was performed following the protocol described by Li *et al*. (2017). For detailed examination of cellular anatomical features, hand-cut cross-sections (∼20 µm thickness) were prepared using a Gillette blade and observed under a Leica DM5000B microscope (Leica, Germany) equipped with a digital imaging system. For lignin staining assay, fresh leaves from tillering-stage were hand-sectioned with a Gillette blade, and then the sections were incubated in 2% (w/v) phloroglucinol-HCl solution for 5 minutes. Stained sections were visualized and observed under Leica DM5000B microscope. Lignin content was determined in leaves of tillering-stage using a commercial assay kit (Beijing Boxbio Science & Technology Co., Ltd., Beijing, China), following the manufacturer’s instructions.

To investigate spatiotemporal expression patterns of *RLC3*, a 2.2-kb genomic DNA fragment upstream of the *RLC3* start codon was amplified from wild-type plants (Nip) and cloned into the binary vector *pCAMBIA1301* to drive GUS reporter gene expression. The recombinant construct (*proRLC3::GUS*) was introduced into Nip callus via *Agrobacterium*-mediated genetic transformation, and T_3_ transgenic plants were selected for subsequent analyses. For GUS histochemical staining, multiple tissues and transverse sections from *proRLC3::GUS* transgenic plants were collected and processed as described by Hu *et al*. (2020) with minor modifications.

### RNA extraction, qRT-PCR and RNA-seq analysis

Total RNA was extracted from rice tissues using RNAiso™ Plus reagent (Takara Bio Inc., Dalian, China) following the manufacturer’s instructions. For first-strand cDNA synthesis, 2 µg of total RNA was reverse transcribed using the Integrated First-strand cDNA Synthesis Kit (with One Step gDNA Remover) from DiNing (Beijing, China). For quantitative real-time PCR (qRT-PCR), reactions were carried out on an Applied Biosystems QuantStudio 5 system with ChamQ Universal SYBR qPCR Master Mix (Vazyme Biotech, Nanjing, China). Gene expression was analyzed via the 2^−ΔΔCT^ method, normalized to the rice UBQ reference gene.

For transcriptome analysis, leaves from Nip and *rlc3-1* plants at the tillering stage were harvested, with each biological replicate consisting of pooled leaf tissue from 5 individual plants. RNA-seq libraries were constructed, sequenced, and subjected to initial data analysis by Beijing Biomarker Technology Co., Ltd., employing standard Illumina protocols. Differentially expressed genes (DEGs) between genotypes were identified through bioinformatics pipelines, including quality control, alignment to reference genomes, and statistical analysis (fold change ≥2, FDR <0.05).

### Subcellular localization assay

To characterize the subcellular localization of RLC3, OSH1, OSH45, and OSH71 proteins, their coding sequences (excluding stop codons) were amplified and cloned into the p35S-eGFP or p35S-mCherry binary vectors under the control of the CaMV 35S promoter. For transient expression assay, the recombinant construct was co-transformed with a nuclear localization marker (NLS-mCherry) into *N. benthamiana* leaves via *Agrobacterium tumefaciens*-mediated infiltration. After 60–72 hours of incubation, leaf epidermal cells were visualized using a confocal laser scanning microscope (Revolution-XD, Andor, UK).

### Protein-protein interaction assay

We used yeast two-hybrid (Y2H) assay, luciferase complementation imaging (LCI) assay, bimolecular fluorescence complementation (BiFC) assay, and GST pull-down assay to characterize the interactions between RLC3 and OSH1, OSH45, OSH71. For Y2H assay, full-length or truncated coding sequences (CDSs) of *RLC3*, *OSH1*, *OSH45*, and *OSH71* were cloned into either pGBKT7 (bait, BD) or pGADT7 (prey, AD) vectors. For interaction screening, various bait-prey combinations were co-transformed into Y2HGold yeast cells following the manufacturer’s protocol (Clontech). For LCI assay, the full-length CDSs of *RLC3* and *OSH1/45/71* were individually fused to the C-terminal fragment (cLUC) or N-terminal fragment (nLUC) of *Renilla luciferase*, respectively, under the control of the CaMV 35S promoter. The recombinant constructs were introduced into *Agrobacterium tumefaciens* strain GV3101 and infiltrated into *N. benthamiana* leaves. After 24 h of dark incubation and 48 h under normal light, leaves were sprayed with 1 mM Beetle luciferin (Promega, E1603) and kept in darkness for 5 min. Luciferase activity was visualized using a low-light CCD imaging system (PlantView100, Biolight Biotechnology). For BiFC assay, the full-length CDSs of *RLC3* and *OSH1/45/71* were cloned into the 2YN (N-terminal YFP fragment) and 2YC (C-terminal YFP fragment) vectors, respectively. The recombinant constructs were transformed into *Agrobacterium* GV3101 and co-expressed in *N. benthamiana* leaves. After 24 h of dark and 48 h of light incubation, YFP fluorescence was detected using a confocal microscope (Revolution-XD, Andor) with 514 nm excitation. For pull-down assay, His-tagged RLC3 (full-length) was expressed from pET-21a(+) in *E. coli*, while GST-tagged OSH1/45/71 were expressed from pGEX-4T-1. Purified His-RLC3 was incubated with glutathione beads pre-bound to GST-OSH1/45/71. After extensive washing, bound proteins were eluted and analyzed by SDS-PAGE/Western blotting using anti-His and anti-GST antibodies (Abmart Inc., Shanghai).

### Transcriptional regulation assays

Yeast one-hybrid (Y1H) assay, dual-luciferase (LUC) assay, and electrophoretic mobility shift assay (EMSA) were used to characterize RLC3 binding to the promoters of *OsPAL5*, *OsCOMT5*, *OsCCR4*, and *OsCAld5H1*/*F5H1*. For Y1H assay, the CDS of RLC3 was cloned into the pB42AD vector to generate the AD-RLC3 fusion construct, while promoter regions of *OsPAL5*, *OsCOMT5*, *OsCCR4*, and *OsCAld5H1* (1.2–2.2 kb upstream of the ATG) were cloned into the pLacZi reporter vector. The recombinant constructs were co-transformed into yeast strain EGY48, transformants were initially cultured on SD/-Ura/-Trp dropout plates for 3 days at 30°C and then transferred to plates containing X-gal to monitor β-galactosidase activity. For dual-LUC assay, the promoter sequences of *OsPAL5*, *OsCOMT5*, *OsCCR4*, and *OsCAld5H1* were cloned into the *pGreenII 0800-LUC* vector to generate reporter constructs, while the full-length CDS of *RLC3* was inserted into the *pGreenII 62-SK* vector as the effector. The recombinant plasmids were transformed into *Agrobacterium tumefaciens* GV3101 (pSoup-p19) and then were co-infiltrated into *N. benthamiana* leaves. After 48–72 hours, luciferase activity (LUC/REN ratio) was quantified using the Dualucif® Firefly & Renilla Assay Kit (Uelandy) on a GloMax 20/20 luminometer (Promega). LUC signals were visualized using a PlantView100 low-light CCD imaging system (Guangzhou Biolight Biotechnology Co., Ltd., Guangzhou, China). For EMSA, a truncated RLC3 coding sequence (amino acids 510–791) was fused in-frame with a GST tag and expressed in *E. coli* using the pGEX4T-1 vector. Biotin-labeled probes (30–50 bp) were incubated with purified recombinant GST-RLC3 protein in EMSA/Gel-Shift Binding Buffer (Beyotime, Shanghai) at room temperature for 30 min. Unlabeled competitors (10×, 25× and 50×) were added for specificity verification. Protein-DNA complexes were resolved on 6% non-denaturing gels in 0.5× TBE at 100 V for 2 h, and then detected with streptavidin-HRP chemiluminescence using ChemiDocXRS+ (Bio-Rad, USA).

### Primer information and Accession numbers

Primer sequences for all applications (genetic mapping, PCR, qRT-PCR, vector construction, and EMSA) are provided in Table S1. Sequence data in this article can be found in the following accession numbers: *RLC3/OsBLH4*, LOC_Os03g52239; *OSH1*, LOC_Os03g51690; *OSH45*, LOC_Os08g19650; *OSH71*, LOC_Os05g03884; *OsPAL5*, LOC_Os04g43760; *OsCOMT5*, LOC_Os04g09654; *OsCCR4*, LOC_Os01g18110; *OsCAld5H1/F5H1*, LOC_Os10g36848.

## Results

### Phenotypic characterization of *rlc3-1 and rlc3-2*

The *rlc3-1* and *rlc3-2* mutants were independently isolated from a mutant library comprising >800 lines. Both mutants displayed adaxial leaf rolling from seedling stage through maturity (Fig. 1a-c), characterized by upward curling of the leaf blade along the adaxial surface (Fig. 1d,e). Quantitative analysis revealed that the leaf rolling index (LRI) of the top three leaves in *rlc3-1* and *rlc3-2* ranged from 16% to 50%, significantly exceeding the LRI of the wild-type Nip ( consistently <3%; Fig. 1f). Beyond leaf rolling, both mutants exhibited pleiotropic alterations, including reduced plant height and leaf width (Fig. 1g,h), with *rlc3-2* additionally showing a significant reduction in leaf length while *rlc3-1* maintained wild-type leaf length (Fig. 1i). Both mutants displayed semi-open glume phenotypes at frequencies of 38.5% (*rlc3-1*) and 11.5% (*rlc3-2*), compared to 1.8% in the wild-type Nip (Table S2). Agronomic analysis revealed that *rlc3-1* exhibited increased grain length, grain width, and 1000-grain weight, whereas *rlc3-2* showed alterations in panicle length, grain length, grain width, and seed setting rate (Table S2), demonstrating distinct yet overlapping impacts on vegetative and reproductive development.

**Fig. 1.**
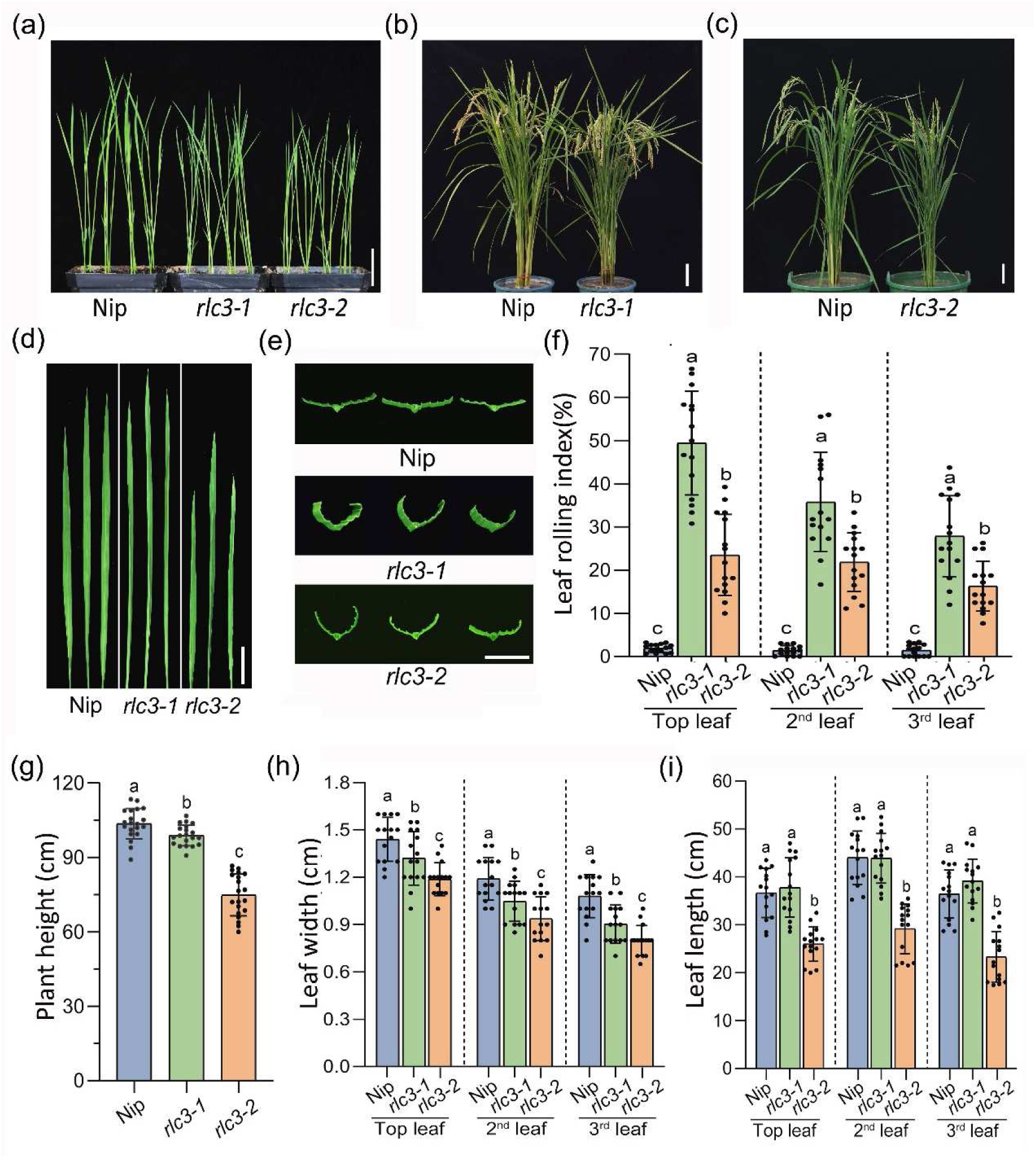
Phenotypic characterization of *rlc3-1* and *rlc3-2* mutants. (a) Two-week-old wild-type (Nip), *rlc3-1* and *rlc3-2* seedlings. (b-c) Phenotypes of *rlc3-1* and *rlc3-2* mutants at the mature stage. (d-e) Morphological comparison of leaf blades (d) and its transverse sections (e) in Nip, *rlc3-1* and *rlc3-2* mutants. (f-i) Comparison of leaf rolling index (f), plant height (g), leaf width (h) and leaf length (i) in Nip, *rlc3-1* and *rlc3-2* at the mature stage. Bars, 3 cm (a), 10 cm (b, c), 5 cm (d) and 1 cm (e). Data are means ± SDs of 15 replicates in (f, h, i) and 20 replicates in (g). Different lowercase letters (a, b, c) indicate significant differences among groups at the *p* < 0.05 level by one-way ANOVA with Tukey’s multiple comparison tests.

### Map-based cloning and functional confirmation of *RLC3*

To elucidate the genetic basis of leaf rolling, *rlc3-1* and *rlc3-2* were crossed with the flat-leaf cultivar Kasalath, respectively. The F_1_ progeny exhibited a uniform flat-leaf phenotype, while the F_2_ population derived from *rlc3-1* × Kasalath showed a 3:1 segregation ratio (443 normal:140 mutant, *χ²* = 0.302, *p* = 0.58 > 0.05), confirming that the leaf rolling phenotype is controlled by a single recessive nuclear gene. Genetic mapping revealed that *rlc3-1* is located on rice chromosome 3, flanked by molecular markers RM15781 and RM85 (Fig. 2a). Further genotyping of 140 F_2_ mutant individuals delimited the gene to a 2.1-cM region between RM15851 and RM3525 (Fig. 2b). Fine-mapping with 429 F^2^ mutant individuals narrowed the candidate interval to a 6-kb physical distance between InDel markers M5/M6/M7 and M8 (Fig. 2c). Genetic mapping confirmed that *rlc3-2* co-segregated with InDel marker R3M30 (Fig. S1), which occupies the same genomic position as M8 on chromosome 3, indicating that *rlc3-1* and *rlc3-2* are allelic variants of the *RLC3* locus. Bioinformatic analysis of the 6-kb genomic interval identified a single annotated gene, LOC_Os03g0732100 (*OsBLH4*), encoding a BELL-like homeodomain protein (Fig. 2d). Sequence analysis revealed distinct mutations: a G>A substitution in the second exon of *rlc3-1* introducing a premature stop codon at 579Trp, and a C>T mutation in the first exon of *rlc3-2* generating a premature stop at 81Gln (Fig. 2d; Fig. S1). These findings demonstrate that *OsBLH4* is the causal gene underlying both *rlc3-1* and *rlc3-2* leaf rolling phenotypes.

**Fig. 2.**
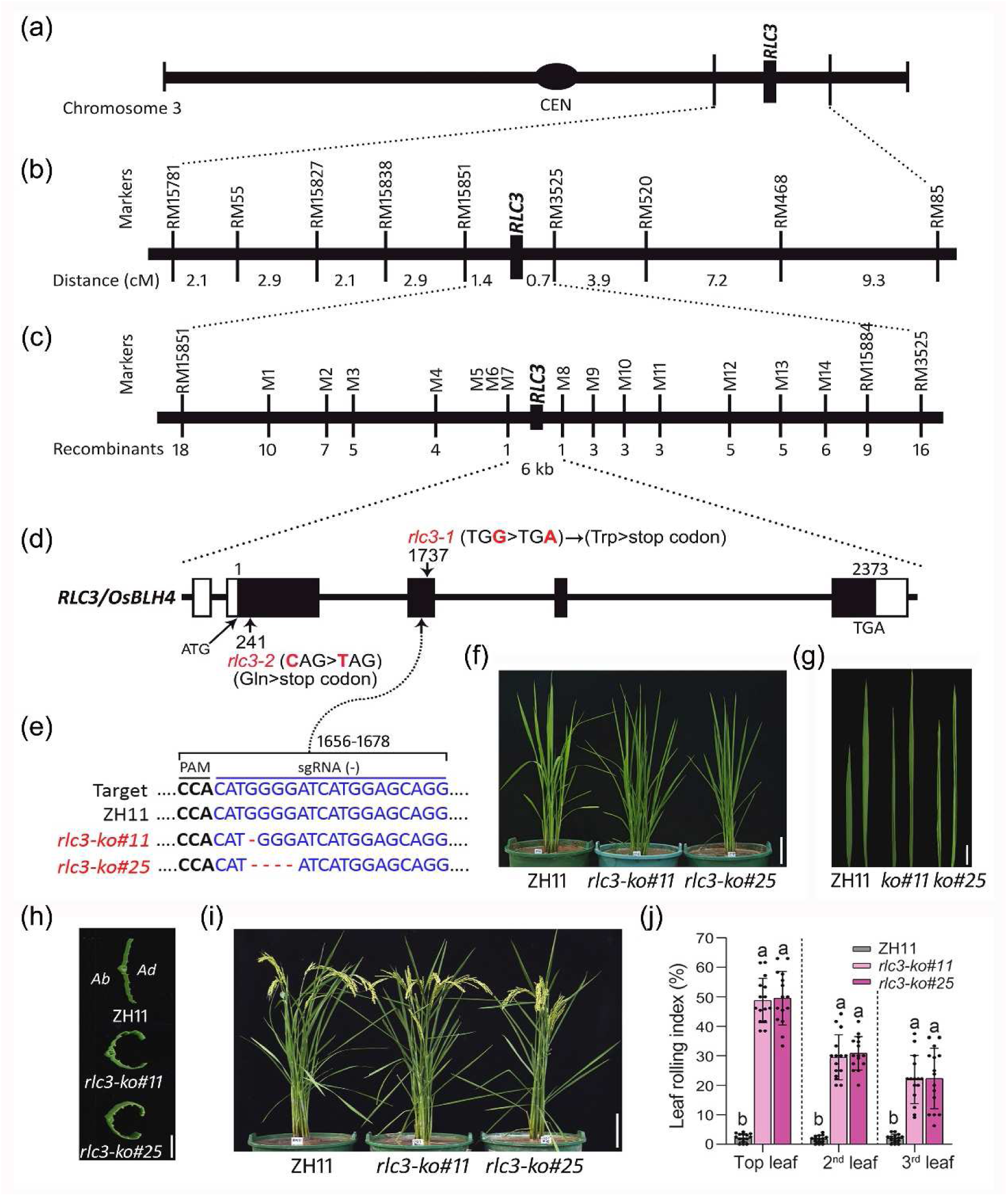
Map-based cloning and functional confirmation of *RLC3*. (a) *RLC3* was genetically mapped to SSR markers on the short arm of chromosome 3 by using an F_2_ segregation population of *rlc3-1*×Kasalath. (b) *RLC3* was further mapped to a 2.1-centimorgan (cM) region flanked by SSR markers RM15851 and RM3525. (c) *RLC3* was fine mapped to a 6-kilobase (kb) physical region flanked by InDel markers M7 and M8. (d) *OsBLH4* (LOC_Os01g52239) is the only candidate gene for *RLC3* within the 6-kb physical interval. The *rlc3-1* and *rlc3-2* mutants carry point mutations in the third and second exons, respectively, both resulting in premature termination of *RLC3/OsBLH4* protein translation. (e) Schematic of CRISPR/Cas9-mediated gene editing in *RLC3/OsBLH4*. Two knockout lines (*rlc3-ko#11* and *rlc3-ko#25*) were identified from over 30 transgenic lines. (f) Morphological comparison of ZH11 (wild-type), *rlc3-ko#11* and *rlc3-ko#25* knock-out plants at the tillering stage. Bar, 10 cm. (g) Phenotypic comparison of the top two leaf blades between ZH11, *rlc3-ko#11* and *rlc3-ko#25* plants at the tillering stage. Bar, 5 cm. (h) Transverse sections of the top leaves in ZH11, *rlc3-ko#11* and *rlc3-ko#25* plants. Bar, 5 mm. (i) Morphological comparison of ZH11, *rlc3-ko#11* and *rlc3-ko#25* plants at the mature stage. Bar, 10 cm. (j) Leaf rolling index of ZH11, *rlc3-ko#11* and *rlc3-ko#25* plants. Data are means ± SDs of 15 replicates; significant differences are indicated by different lowercase letters according to one-way ANOVA with Tukey’s multiple comparison tests.

To functionally validate *RLC3*, CRISPR/Cas9-mediated knockout lines targeting *OsBLH4* were generated in rice Zhonghua 11 (ZH11). Two homozygous mutants (*rlc3-ko#11* and *rlc3-ko#25*) were identified, harboring 1-bp and 4-bp deletions, respectively, both causing frameshift mutations in the third exon (Fig. 2e). Phenotypic analysis revealed consistent leaf rolling phenotypes from seedling stage through maturity in both knockout lines (Fig. 2f-i), phenocopying the *rlc3-1* and *rlc3-2* mutant phenotypes. Quantitative assessment of leaf rolling index (LRI) demonstrated substantial increases in the top three leaves (50%, 30%, and 20% for knockout lines) compared to wild-type plants with near-zero LRI values (Fig. 2j). These results confirm that *OsBLH4* is the functional ortholog of *RLC3*, with loss-of-function mutations causing the leaf rolling phenotype.

To conclusively demonstrate that all *rlc3* mutant lines result from *RLC3/OsBLH4* mutations, reciprocal crosses and allelism tests were performed between *rlc3-1*, *rlc3-2*, and *rlc3-ko#25*. F_1_ hybrids from *rlc3-1* × *rlc3-2* and *rlc3-1* × *rlc3-ko#25* crosses were confirmed as heterozygotes by DNA sequencing (Fig. S2). These F_1_ plants exhibited consistent leaf rolling phenotypes with LRI values comparable to *rlc3-1*, *rlc3-2*, and *rlc3-ko#25* (Fig. S3). These results provide definitive genetic evidence that *rlc3-1*, *rlc3-2*, and *rlc3-ko#25* are fully allelic mutations of *RLC3*, confirming *OsBLH4* (*LOC_Os03g52239*) as the causal gene.

### RLC3, a nuclear-localized BELL-type homeobox transcription factor, is constitutively expressed in rice

Bioinformatic analysis and RT-PCR validation revealed that the *RLC3/OsBLH4* locus generates three alternatively spliced mRNA transcripts. The longest isoform (LOC_Os03g52239.1, herein designated RLC3) encodes a 791 aa protein (81.9 kDa, pI 6.95) containing POX (aa 373-509) and Homeobox (aa 581-618) domains (Fig. S4). The two shorter isoforms—LOC_Os03g52239.2 (*RLC3-v2*) and LOC_Os03g52239.3 (*RLC3-v3*)—exhibit truncated CDS (1026 bp and 966 bp, respectively) and shifted domain locations (Fig. S4). Comparative 3D modeling via SWISS-MODEL demonstrated distinct structural architectures among isoforms (Fig. S4). Given its intact domain organization, *RLC3* (LOC_Os03g52239.1) was selected as the primary study subject unless otherwise specified.

RLC3/OsBLH4 protein shares 37.13% and 35.18% similarity to Arabidopsis AtBLH2/SAW1 and AtBLH4/SAW2. Multiple sequence alignments revealed high conservation of both POX and Homeobox domains between RLC3 and its Arabidopsis homologs AtBLH2/AtBLH4, with the Homeobox domain showing particularly strong conservation (Fig. S5), suggesting shared molecular functions. Phylogenetic analysis demonstrated distinct evolutionary clades between monocots and dicots (Fig. S5). Subcellular localization assay revealed that RLC3-GFP fluorescence exclusively colocalized with the nuclear marker (Fig. 3a), confirming its nuclear localization. The two shorter isoforms (RLC3-v2 and RLC3-v3) also demonstrated co-localization with nuclear marker (Fig. S6). To test RLC3’s transcriptional self-activation, its full-length CDS was cloned into pGBKT7 and transformed it into Y2HGold yeast cells. Growth on SD/-Trp-His-Ade medium with X-α-Gal confirmed self-activation activity (Fig. 3c). To map the functional domains, we generated ten RLC3 truncations based on its conserved POX (373-509 aa) and Homeobox (581-618 aa) domains (Fig. 3b). Among these, the N-terminus (1-372 aa) and C-terminus (619-791 aa) constructs enabled yeast growth on SD/-Trp-His-Ade and induced blue pigmentation with X-α-Gal (Fig. 3c), suggesting these regions mediate self-activation. Combined with its nuclear localization, these results demonstrate that RLC3 functions as a nuclear transcription factor.

**Fig. 3.**
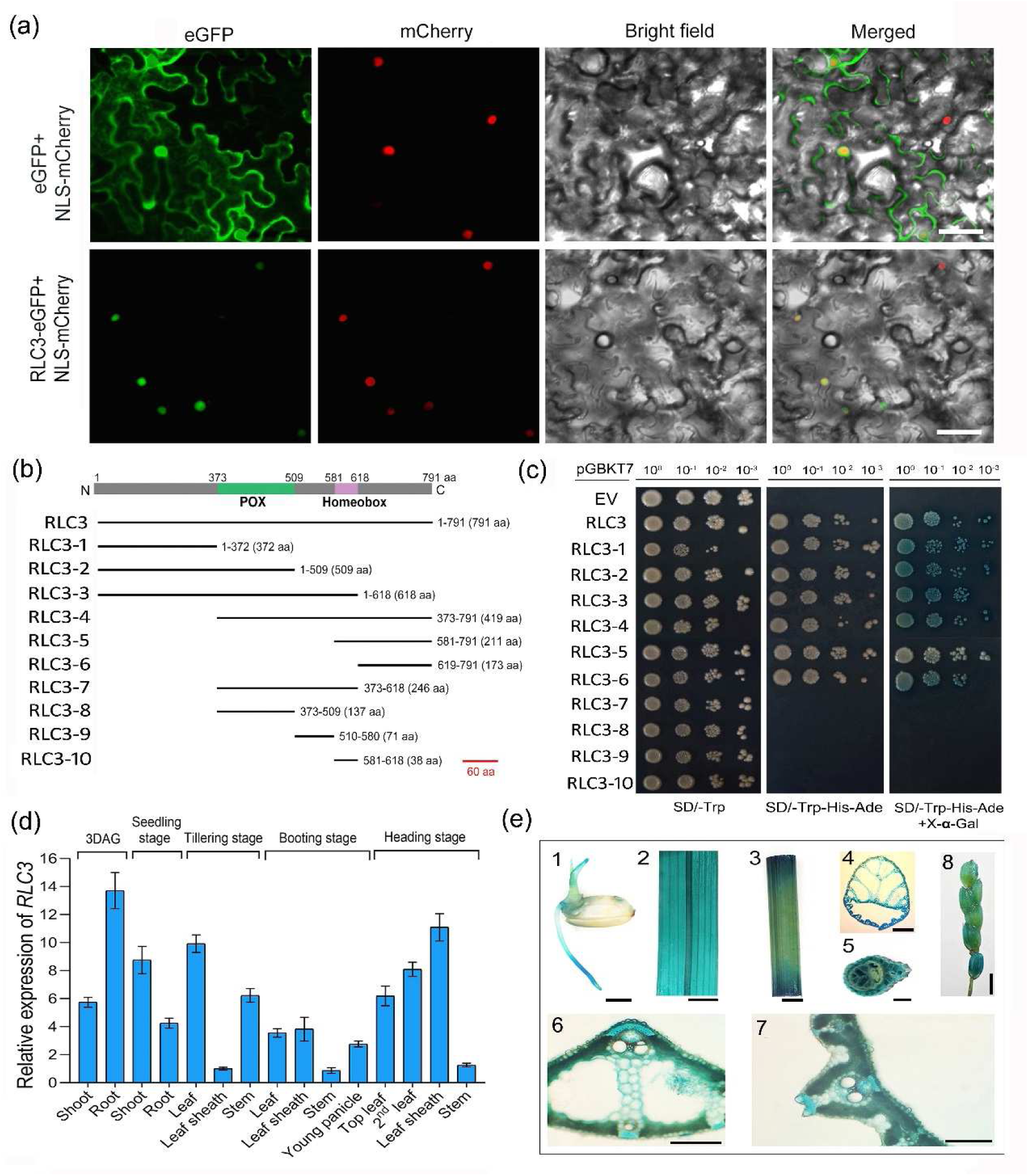
Expression analysis of RLC3. (a) Subcellular localization of RLC3 protein in *Nicotiana benthamiana*. RLC3-eGFP and NLS-mCherry constructs (nuclear localization marker) were co-infiltrated into leaf epidermal cells of *Nicotiana benthamiana* via Agrobacterium-mediated transformation. The infiltrated leaf epidermal cells were then observed using a confocal laser scanning microscope with 488 nm excitation for GFP detection (green fluorescence) and 561 nm excitation for mCherry detection (red fluorescence). Bars, 50 µm. (b) Domain architecture schematics of full-length and truncated RLC3 constructs. N, N-terminus; C, C-terminus. (c) Autoactivation assessment of full-length and truncated RLC3 variants in yeast two-hybrid (Y2H) system. EV, empty vector pGBKT7. Numbers above panels indicate dilution factors (10^0^∼10^-3^). (d) Spatiotemporal expression profiling of *RLC3* in rice tissues by qRT-PCR. Expression levels were normalized to the *Ubiquitin* (*Ubq*) internal reference gene, with transcript levels in tillering-stage leaf sheaths set as 1.0 (calibrator). Data are means ± SDs of three replicates. (e) Histochemical localization of *RLC3* promoter-driven GUS activity. Representative staining patterns in transgenic plants are shown: 3-day-old germinated seeds (e1), the second leaf (e2), stems (e3), cross-sections of leaf sheath (e4), stem (e5), leaf midrib (e7) and leaf vein (e8) at the tillering stage, and young panicles at the heading stage (e6). Bars, 2 mm (e1, e4, e5), 5 mm (e2, e3, e8) and 500µm (e6, e7).

RT-qPCR demonstrated RLC3’s ubiquitous expression across rice developmental stages, with peak levels in tillering-stage leaves (Fig. 3d), correlating with the mutant’s leaf-rolling phenotype. *RLC3* promoter-GUS staining confirmed expression in germinating seeds, tillering-stage leaves/stems, and young panicles (Fig. 3e), consistent with qRT-PCR data. Detailed anatomical analysis revealed GUS signals in mesophyll, parenchyma, vascular bundles, and mechanical tissues of leaves, as well as stem parenchyma and vascular tissues (Fig. 3e), supporting *RLC3*’s role in leaf morphogenesis. Spatiotemporal expression analysis of *RLC3-v2* (LOC_Os03g52239.2) and *RLC3-v3* (LOC_Os03g52239.3) transcripts revealed patterns similar to those of *RLC3* (Fig. S6).

### Loss-of-function mutations in *RLC3* disrupt bulliform cells development and midrib formation

To investigate the cytological basis of leaf rolling, we analyzed paraffin cross-sections of WT and *rlc3* mutant leaves. Microscopic observation revealed that both *rlc3-1* and *rlc3-2* mutants exhibited significantly smaller bulliform cells (BCs) compared to Nip (Fig. 4a,c; Fig. S7). Quantitative analysis showed *rlc3-1* and *rlc3-2* have 25.7% and 21.5% reduction in BC area adjacent to large veins (LVs), and a 30.1% and 27.5% reduction between small veins (SVs), respectively (Fig. 4e). Notably, *rlc3-1* displayed a more severe BC area reduction between SVs (3.5% greater than *rlc3-2*), correlating with its stronger leaf rolling phenotype. Additionally, the relative depth of BCs (BCs depth/leaf thickness ratio) was significantly reduced in both mutants (Fig. 4f), consistent across all vein positions. All these defects were further confirmed in knockout lines *rlc3-ko#11* and *rlc3-ko#25* (Fig. 4b, d, g, h; Fig. S7), demonstrating that *RLC3* loss-of-function disrupts BCs development.

**Fig. 4.**
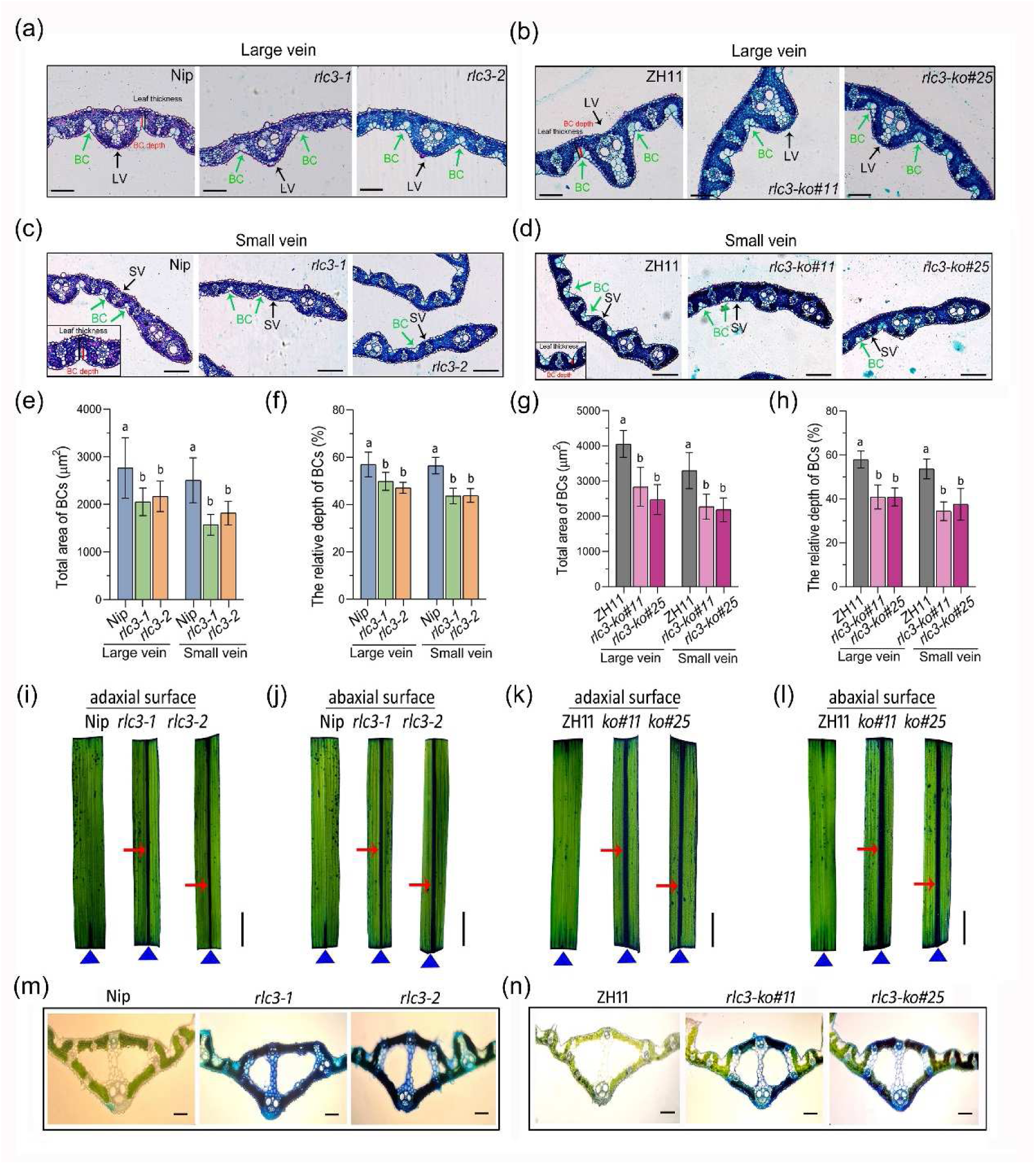
Cytological and histological analysis of *RLC3* mutants. (a-d) Cross-sections of wild-type and mutant leaves. The middle region of the second leaf blade from tillering-stage plants was used for paraffin sectioning. (a-b) show the area near the midrib, while (c-d) show the area near the leaf margin. BC, bulliform cell; LV, large vein; SV, small vein. Bars, 100 µm. (e, g) BCs area quantification. BCs adjacent to LVs or between SVs were analyzed separately. (f, h) BCs relative depth analysis. BCs relative depth is calculated as the ratio of BCs depth to leaf thickness. Data in (e-h) are means ± SDs of 30 replicates. Significant differences are indicated by different lowercase letters according to one-way ANOVA with Tukey’s multiple comparison tests. (i-l) Toluidine blue (TB) staining of leaf blades with both ends removed. The adaxial (upper) and abaxial (lower) surfaces of the leaf are shown, respectively. The position of the midrib is indicated by triangle, and the stained midrib is indicated by arrow. Bars, 2 cm. (m-n) Transverse sections of TB-stained leaf blades, showing midrib positions. Bars, 300 µm.

Furthermore, we employed toluidine blue (TB) staining on leaf blades with both ends removed. A striking difference was observed in the midrib: the ends-removed wild-type leaves completely repelled TB staining, whereas all four mutant lines (*rlc3-1*, *rlc3-2*, *rlc3-ko#11*, and *rlc3-ko#25*) exhibited obvious TB staining at midrib (Fig. 4i-l), indicating severe midrib defects. Free-hand cross sections were further prepared from TB-stained leaves and examined under microscopy. Consistent with the TB staining results, the midribs of wild-type (Nip and ZH11) exhibited minimal TB staining, whereas all four mutant lines (*rlc3-1*, *rlc3-2*, *rlc3-ko#11*, and *rlc3-ko#25*) showed intense staining throughout the entire midrib (Fig. 4m-n). As illustrated in cross sections, the mutant lines displayed extensive staining in sclerenchyma cells and midrib vascular tissues, including xylem, phloem, and mechanical tissues (Fig. 4m-n). Additionally, pronounced staining was observed in both large and small veins adjacent to the midrib (Fig. 4m-n). These results collectively indicate that the *RLC3* mutant lines exhibit severe defects in midrib vascular tissues. Together, these findings demonstrate that *RLC3* plays a critical role in the development of BCs and leaf midrib vascular tissues, whose defects leading to leaf rolling.

### Loss-of-function of *RLC3* accelerates water loss in leaves, leading to significantly reduced drought tolerance

Given that the loss-of-function in *RLC3* causes severe defects in leaf midrib and BCs, we further investigated leaf water transport under non-stressed conditions. The relative water content (RWC) was significantly reduced in leaves of all four mutant lines (*rlc3-1*, *rlc3-2*, *rlc3-ko#11*, and *rlc3-ko#25*) compared to wild-type plants (Nip and ZH11) (Fig. 5a). The rate of water loss (RWL) from detached leaves was significantly higher in the four mutant lines than in wild-type at corresponding time points (Fig. 5b). These findings demonstrate that *RLC3* loss-of-function accelerates water loss in leaves.

**Fig.5.**
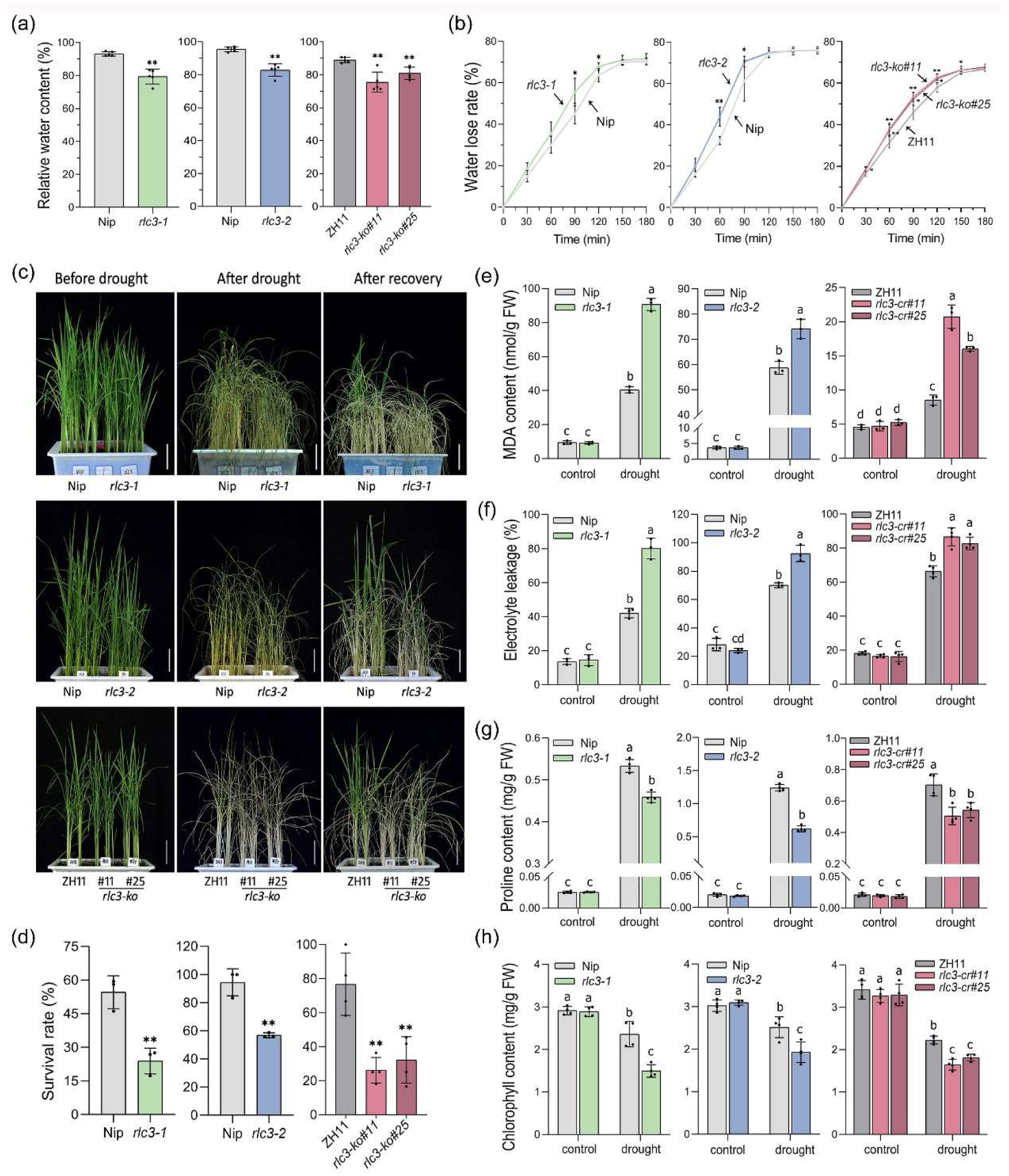
Evaluation of drought stress resistance and water physiological analysis in *RLC3* mutants. (a-b) Comparison of relative water content (a) and water loss rate (b) in leaves of wild-type (Nip and ZH11) and *RLC3* mutant lines (*rlc3-1*, *rlc3-2*, *rlc3-ko#11* and *rlc3-ko#25*). Data are means ± SDs of five biological replicates; asterisks indicate significant differences between the mutant and its wild-type according to Student’s t-test (\*\**P* < 0.01). (c) Drought tolerance of WT and *RLC3* mutant plants. Phenotypes of wild-type and mutant lines before drought stress, after drought stress, and after recovery are shown. Bars, 8 cm. (d) Survival rates of wild-type and *RLC3* mutants. Data are means ± SDs from three independent experiments in *rlc3-1* and *rlc3-2*, respectively, and four independent experiments in *rlc3-ko#11* and *rlc3-ko#25* lines; asterisks indicate significant differences between the mutant and its wild-type according to Student’s t-test (\*\**P* < 0.01). (e-h) Measurement of MDA content (e), electrolyte leakage rate (f), proline content (g), and chlorophyll content (h) in leaves of wild-type and *RLC3* mutant lines under non-stress (control) and drought stress conditions. Data are means ± SDs from four biological replicates; different lowercase letters (a, b, c) indicate significant differences among groups at the *p* < 0.05 level by one-way ANOVA with Tukey’s multiple comparison tests.

We then compared drought tolerance between wild-type and the mutant lines. At the seedling stage, all four mutant lines (*rlc3-1*, *rlc3-2*, *rlc3-ko#11*, and *rlc3-ko#25*) exhibited significantly reduced drought tolerance compared to wild-type (Fig. 5c-d), and similar results were observed at the grain-filling stage (Fig. S8). Under drought stress conditions, all four mutant lines showed significantly higher malondialdehyde (MDA) contents and electrolyte leakage (Fig. 5e-f), along with lower proline and chlorophyll levels compared to wild-type plants (Fig. 5g-h). These findings collectively demonstrate that *RLC3* loss-of-function accelerates water loss and markedly increases drought sensitivity.

### RLC3 is a key regulator of lignin biosynthesis and secondary cell wall formation in leaves

To elucidate the molecular function of *RLC3*, we conducted comparative transcriptome analysis between wild-type (Nip) and *rlc3-1*, identifying 3,750 DEGs (1,342 up-/2,408 down-regulated) (Table S3). Gene Ontology (GO) enrichment analysis showed DEGs were primarily enriched in catalytic activity/binding (molecular function) and metabolic processes (biological process) (Fig. S9). KEGG enrichment analysis revealed phenylpropanoid biosynthesis (ko00940) as the most enriched pathway, with predominant down-regulation, and the related pathways—phenylalanine/tyrosine/tryptophan (ko00400) and flavonoid biosynthesis (ko00941)—were also significant enriched (Fig. 6a; Fig. S10-S12). Notably, more than 30 DEGs in phenylpropanoid biosynthesis pathway were annotated as lignin biosynthesis genes (Fig. 6b; Table S4). The lignin biosynthesis pathway involves key intermediates including cinnamic acid, p-coumaric acid, coumaroyl-CoA, flavonoids, and various aldehydes. Key enzymes such as phenylalanine ammonialyase (PAL), cinnamic acid 4-hydroxylase(C4H), 4-coumarate-coenzyme A ligase (4CL), coumarate 3-hydroxylase (C3H), caffeoyl coenzyme A 3-O-metyltransferase (CCoAoMT), cinnamoyl-CoA reductase (CCR), ferulate 5-hydroxylase (F5H), caffeic acid O-methyltransferase (COMT) and cinnamyl alcohol dehydrogenase (CAD) catalyze these lignin biosynthetic reactions (Weng and Chapple, 2010). In this study, transcriptomic analysis identified 31 DEGs (5 up-regulated and 26 down-regulated) associated with lignin biosynthesis pathway, which can be clustered into 11 groups (Fig. 6b). In these lignin-related DEGs, five PAL genes (Tonnessen *et al*., 2015), six CCR genes (Park *et al*., 2017), four COMT genes (Liang *et al*., 2022), three 4CL-encoding genes, three F3H (flavanone 3’-hydroxylase) genes and two CAD genes exhibited down-regulated expression in *rlc3-1* (Fig. 6b). *F5H1/OsCAld5H1*, a major factor controlling S/G lignin composition (Takeda *et al*., 2017), also down-regulated in *rlc3-1* (Fig. 6b). Overall, these data indicate that *RLC3* loss-of-function significantly inhibits gene expression in lignin biosynthesis pathway.

**Fig.6.**
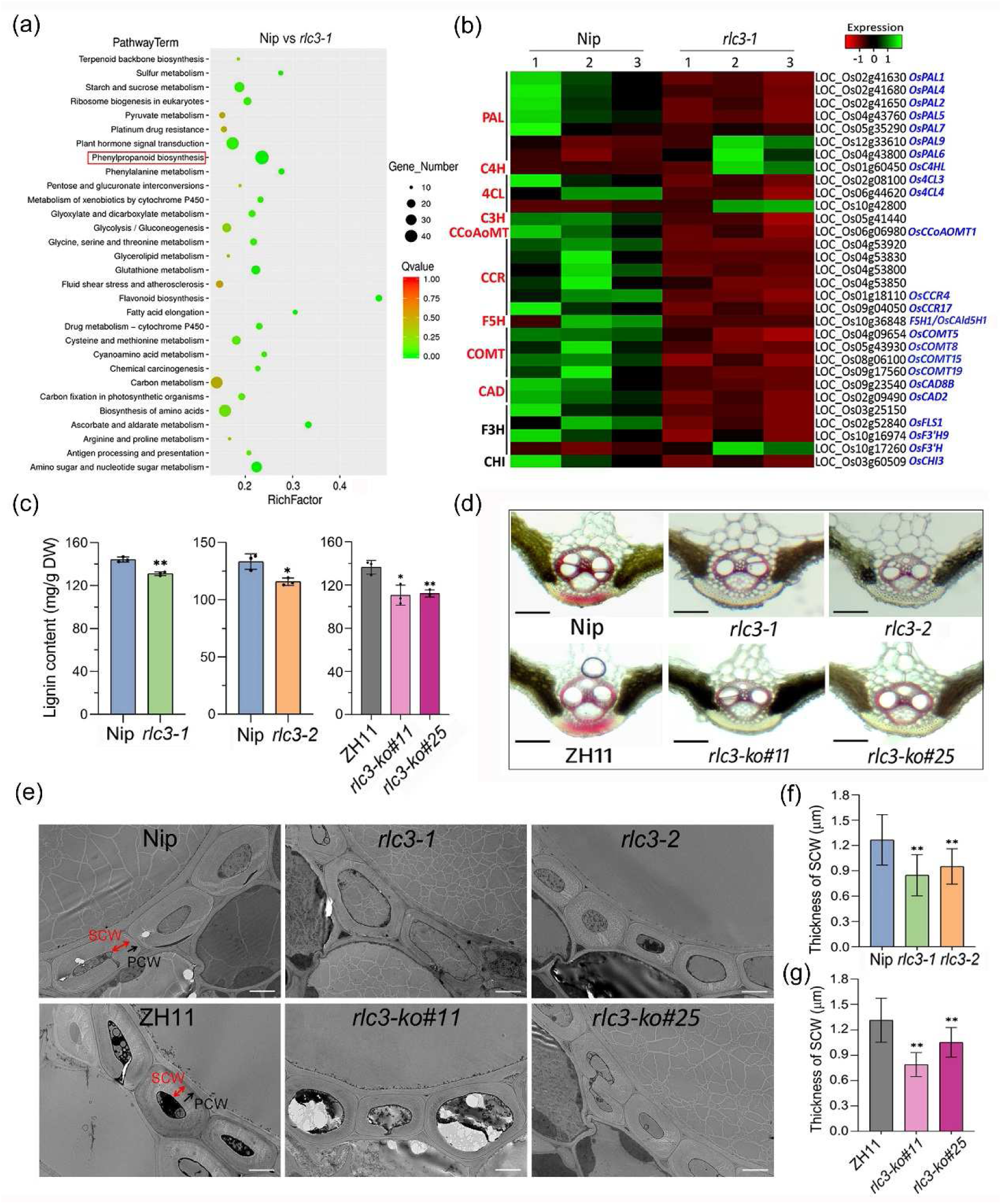
*RLC3* regulates lignin biosynthesis and secondary cell wall formation. (a) KEGG pathway enrichment for differentially expressed genes (DEGs) between wild-type (Nip) and *rlc3-1* mutants. The size of each dot represents the number of DEGs in the pathway, and the color corresponds to different Q-value ranges. (b) Expression heatmap of lignin synthesis-related genes selected from the total DEGs. The green indicating upregulation and red indicating downregulation. (c) Determination of lignin content in leaves of wild-type (Nip and ZH11) and *RLC3* mutant lines (*rlc3-1*, *rlc3-2*, *rlc3-ko#11* and *rlc3-ko#25*). Data are means ± SDs of four biological replicates; asterisks indicate significant differences between the mutant and its wild-type according to Student’s t-test (\**P* < 0.05; \*\**P* < 0.01). (d) Histochemical analysis of lignin in cross sections of leaf midrib vascular bundle using phloroglucinol-HCl staining. Phloroglucinol-HCl stained the walls of xylem cells in the wild-type, whereas weak staining was observed in *RLC3* mutants. Scale bars = 500 µm. (**e**) Transmission electron microscopy observation of cell wall structure in the midrib vascular bundle region of wild-type and mutant leaves. PCW, primary cell wall; SCW, secondary cell wall. Bars = 2 µm. (f-g) Statistics of secondary cell wall thickness in the midrib vascular bundle region of leaves. Data are means ± SD of 30 replicates; asterisks denote significant differences (Student’s t-test, \*\**P* < 0.01).

Furthermore, field-grown leaf samples from *rlc3-1*, *rlc3-2*, and knockout lines (*rlc3-ko#11*, *rlc3-ko#25*) showed significant reductions in lignin content (Fig. 6c), suggesting impaired lignin biosynthesis or altered metabolic pathways. Phloroglucinol-HCl staining of lignin in leaf midribs further confirmed these findings: WT vascular bundles and mechanical tissues exhibited characteristic deep pink staining, whereas mutant lines showed markedly reduced staining intensity, particularly in mechanical tissue regions (Fig. 6d). We then prepared ultrathin sections and observed the ultrastructure of vascular bundles in the midrib region using TEM. It revealed a significant reduction in the secondary cell wall (SCW) thickness of vascular bundles in both *rlc3-1* and *rlc3-2* mutants compared to Nip (Fig. 6e-f). This phenotype was further corroborated in the *rlc3-ko#11*, *rlc3-ko#25* (Fig. 6e). Similarly, when compared to ZH11, the *rlc3-ko#11* and *rlc3-ko#25* lines exhibited diminished SCW thickness in vascular bundles (Fig. 6g). These findings collectively demonstrate that *RLC3* loss-of-function negatively impacts secondary cell wall formation in leaf midribs. Together, our results suggest that *RLC3* plays an essential regulatory role in lignin biosynthesis and secondary cell wall formation.

### RLC3 interacts with KNOX proteins (OSH1/45/71) to coordinately regulate lignin biosynthesis and secondary cell wall formation

To elucidate the regulatory network of RLC3, we identified its interacting partners via a combination of experimental assays. Comprehensive searches in PRIN databases revealed that RLC3 potentially interacts with KNOX members OSH1, OSH45, and OSH71 (Fig. S13). This prediction aligns well with the evolutionarily conserved BELL-KNOX interaction framework observed in several plant species (Tsuda *et al*., 2017; Yoon *et al*., 2017; Tao *et al*., 2018; Wu *et al*., 2025). To verify these interactions, we conducted yeast two-hybrid (Y2H) assay using a RLC3 truncated fragment (373-618 aa) encompassing the conserved POX and Homeobox domains, which were selected based on autoactivation test results (Fig. 3b-c). The Y2H results demonstrated interactions between RLC3 and all three OSH proteins (OSH1, OSH45 and OSH71) (Fig. 7a). Domain mapping experiments further revealed that the conserved POX domain (373-509 aa) of RLC3 specifically mediates these interactions (Fig. 7b).

**Fig.7.**
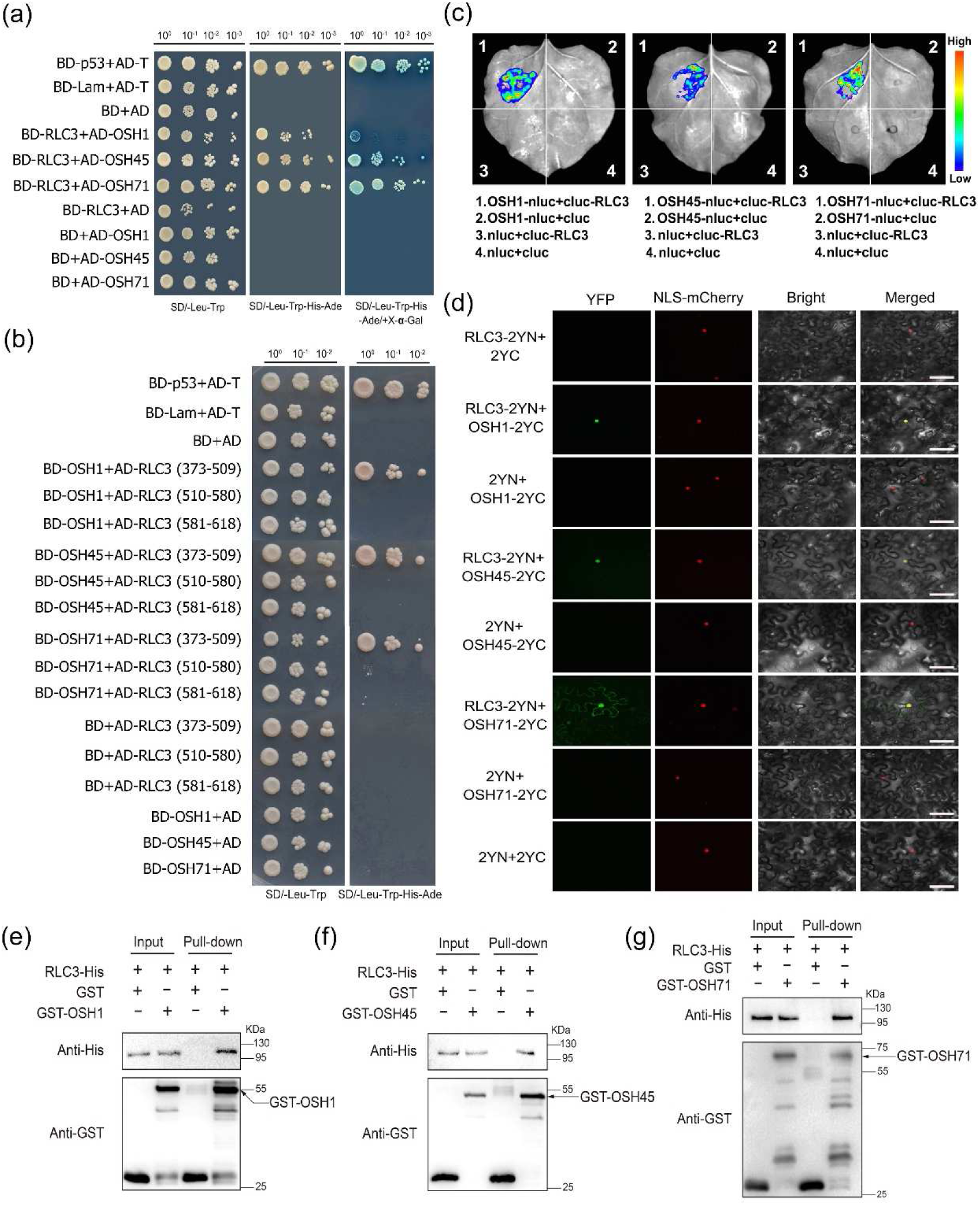
RLC3 interacts with OSH1, OSH45 and OSH71 both in vivo and in vitro. (a) Interaction between RLC3 and OSH1/45/71 in yeast. Y2H analysis was performed to detect the interaction between RLC3 and OSH1/45/71, using BD-p53+AD-T as the positive control and BD-Lam+AD-T as the negative control. BD+AD served as an empty vector negative control. The top number in the figure indicates the yeast dilution. (b) Analysis of interaction domain between RLC3 and OSH1/45/71. Y2H experiment was conducted to detect the interaction between truncated RLC3 and OSH1/45/71. The numbers in the figure represent the amino acid regions of truncated RLC3 proteins. (c) Luciferase complementation imaging (LCI) assay demonstrating the in vivo interaction between RLC3 and OSH1/45/71. (d) Bimolecular fluorescence complementation (BiFC) assay confirming the in vivo interaction between RLC3 and OSH1/45/71. Bars, 50 µm. (e-g) Pull-down assay demonstrating the in vitro physical interactions between RLC3 and OSH1 (e), OSH45 (f), and OSH71 (g) proteins. Recombinant GST-OSH1/45/71 were incubated in binding buffer containing glutathione-agarose beads with or without RLC3-His, respectively. Input and eluted pull-down fractions were analyzed via SDS-PAGE followed by Western blotting with the respective antibodies.

The interactions between RLC3 and OSH1/45/71 proteins were then validated through both LCI assay and BiFC assay (Fig. 7c-d). Moreover, the subcellular co-localization analysis revealed that RLC3-eGFP completely overlapped with OSH1/45-mCherry signals in the nucleus (Fig. S14), while with OSH71-mCherry, significant nuclear co-localization was accompanied by weak membrane-associated co-localization (Fig. S14). This aligns with the BiFC observation where OSH71 partially relocalized RLC3 out of the nucleus (Fig. 7d), suggesting OSH71 may regulate RLC3’s dimer function by modulating its subcellular distribution. Furthermore, GST pull-down assay demonstrated direct physical interaction between RLC3 and OSH1/44/71 in vitro (Fig. 7e-g). These results suggest that RLC3 forms functional BELL-KNOX complexes with OSH1, OSH45, and OSH71 in the nucleus.

To functionally characterize the RLC3-OSH1/45/71 interactions, we generated knock-out lines of OSHs using CRISPR/Cas9-mediated gene editing (Fig. 8a-c). Phenotypic analysis revealed distinct morphological alterations in *OSH1* and *OSH45* mutants compared to wild-type plants: both *osh1-ko* and *osh45-ko* exhibited mild leaf rolling, reduced plant height, and altered leaf length and width (Fig. 8d-i; Fig. S15). In contrast, *osh71-ko* mutants showed flat leaves, reduced plant height, and shorter leaf length relative to wild-type plants (Fig. 8j-l; Fig. S15).

**Fig.8.**
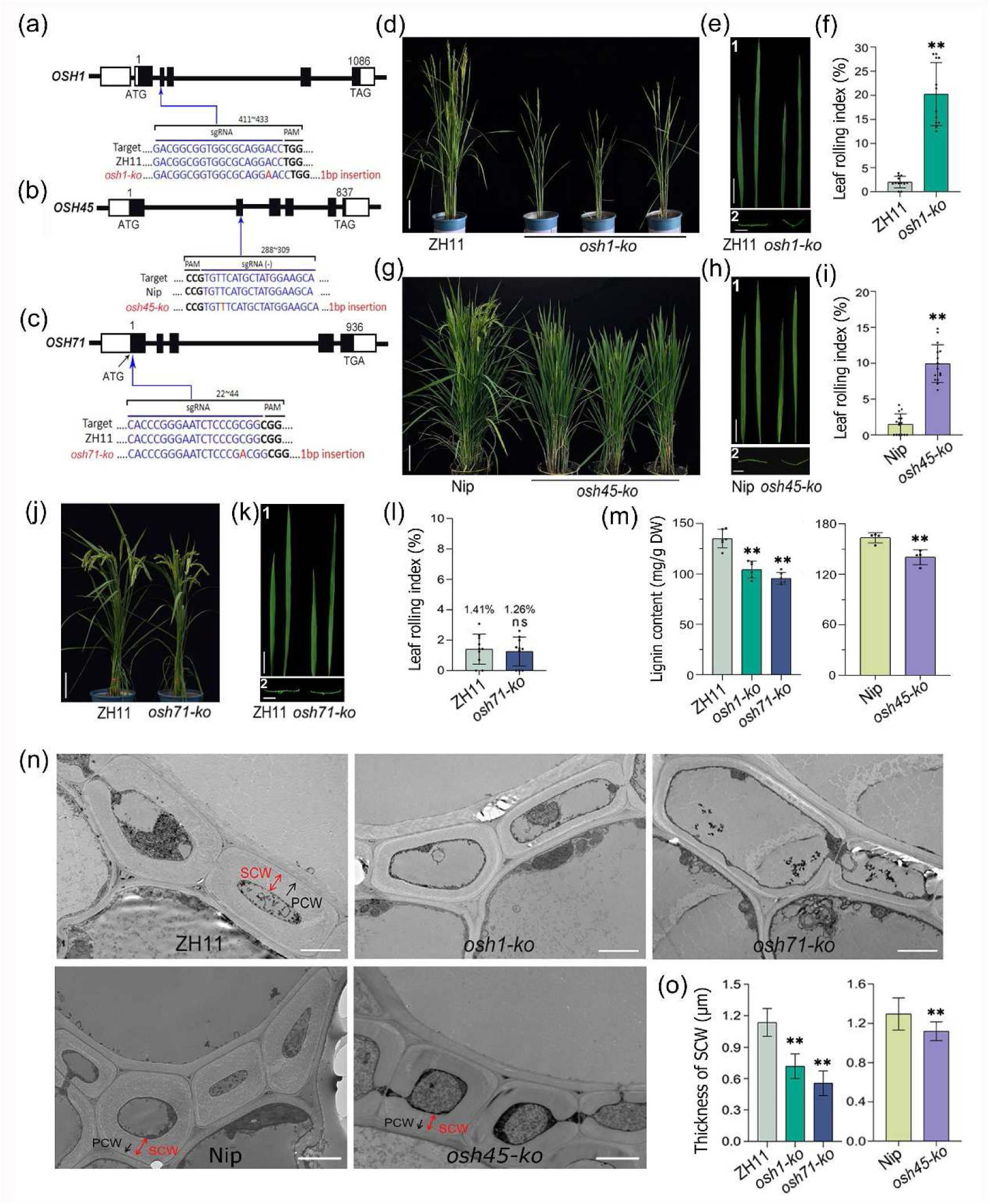
Loss-of-function of *OSH1/45/71* disrupts lignin biosynthesis and secondary cell wall formation, leading to altered leaf morphology. (a-c) Schematic of CRISPR/Cas9-mediated knockout of *OSH1*, *OSH45*, and *OSH71*. (d-f) Plant morphology (d), leaf blade (e1) and transverse section (**e2**), and leaf rolling index statistics (f) of mature-stage wild-type ZH11 and *osh1-ko* mutant. (g-i) Plant morphology (g), leaf blade (h1) and transverse section (h2), and leaf rolling index statistics (i) of mature-stage wild-type Nip and *osh45-ko* mutant. (j-l) Plant morphology (j), leaf blade (k1) and transverse section (k2), and leaf rolling index statistics (l) of mature-stage wild-type ZH11 and *osh71-ko* mutant. Data are means ± SDs (n=10); asterisks indicate significant differences between the mutant and its wild-type according to Student’s t-test (\*\**P* < 0.01; *ns*, no significant difference). Bars, 20 cm in (d, g, j), 8 cm in (e1, h1, k1), 5 mm in (e2, h2, k2). (m) Quantification of lignin content in leaves of wild-type (Nip and ZH11) and osh1-ko, *osh45-ko* and *osh71-ko* mutants. Data are means ± SDs from four biological replicates; asterisks indicate significant differences between the mutant and its wild-type according to Student’s t-test (\*\**P* < 0.01). (n) Transmission electron microscopy observation of cell wall structure in the leaf midrib vascular bundle region of wild-type (Nip and ZH11) and osh1-ko, *osh45-ko* and *osh71-ko* mutants. *PCW*: primary cell wall; *SCW*: secondary cell wall. Bars, 2 µm. (o) Statistics of secondary cell wall (SCW) thickness in wild-type (Nip and ZH11) and *osh1-ko*, *osh45-ko* and *osh71-ko* mutants. Data are means ± SD from 30 replicates; asterisks denote significant differences (Student’s t-test, \*\**P* < 0.01).

To further investigate the role of OSHs in cell wall development, we conducted ultrastructural analysis of leaf midribs to compare xylem cell wall architecture between OSHs knock-out lines and the WT control (Fig. 8o). Comparative transmission electron microscopy (TEM) demonstrated a marked reduction in xylem cell wall thickness across all *osh1-ko*, *osh45-ko*, and *osh71-ko* mutants relative to WT (Fig. 8p). Accompanying this structural modification, substantial decreases in lignin content were also observed in *osh1-ko*, *osh45-ko*, and *osh71-ko* leaves (Fig. 8m-n). Collectively, these findings indicate that OSH1, OSH45, and OSH71 are essential for regulating xylem cell wall formation and lignin deposition, by interacting with RLC3.

### RLC3 directly binds to the promoters of lignin biosynthetic genes *OsPAL5*, *OsCOMT5*, *OsCCR4*, and *OsCAld5H1*/*F5H1*and activates their transcription

To validate the regulatory role of RLC3 in lignin biosynthesis, yeast one-hybrid (Y1H) screening was conducted using 31 lignin-related gene promoters. The results demonstrated that RLC3 specifically binds to the promoters of *OsPAL5*, *OsCOMT5*, *OsCCR4*, and *OsCAld5H1*/*F5H1* (Fig. 9a). In vivo binding of RLC3 to the four lignin genes was further demonstrated using a dual-luciferase (LUC) reporter system (Fig. 9b-f). *RLC3* was cloned and inserted into the pGreenII 62-SK vector as the effector, and the promoters of the lignin genes were cloned and inserted into the pGreenII 0800-LUC vector as the reporter gene (Fig. 9b). When RLC3 was co-transformed with each of the four promoters, LUC activity was significantly greater than that in the control (Fig. 9c-f), indicating that RLC3 can enhance the expression of the *LUC* gene by binding to the promoters of *OsPAL5*, *OsCOMT5*, *OsCCR4*, and *OsCAld5H1*/*F5H1*. Additionally, qRT-PCR analysis confirmed that *OsPAL5*, *OsCOMT5*, *OsCCR4*, and *OsCAld5H1*/*F5H1* gene expression were significantly down-regulated in *rlc3* mutants (Fig. S16), indicating that RLC3 positively regulates the transcriptional expression of these lignin genes.

**Fig. 9.**
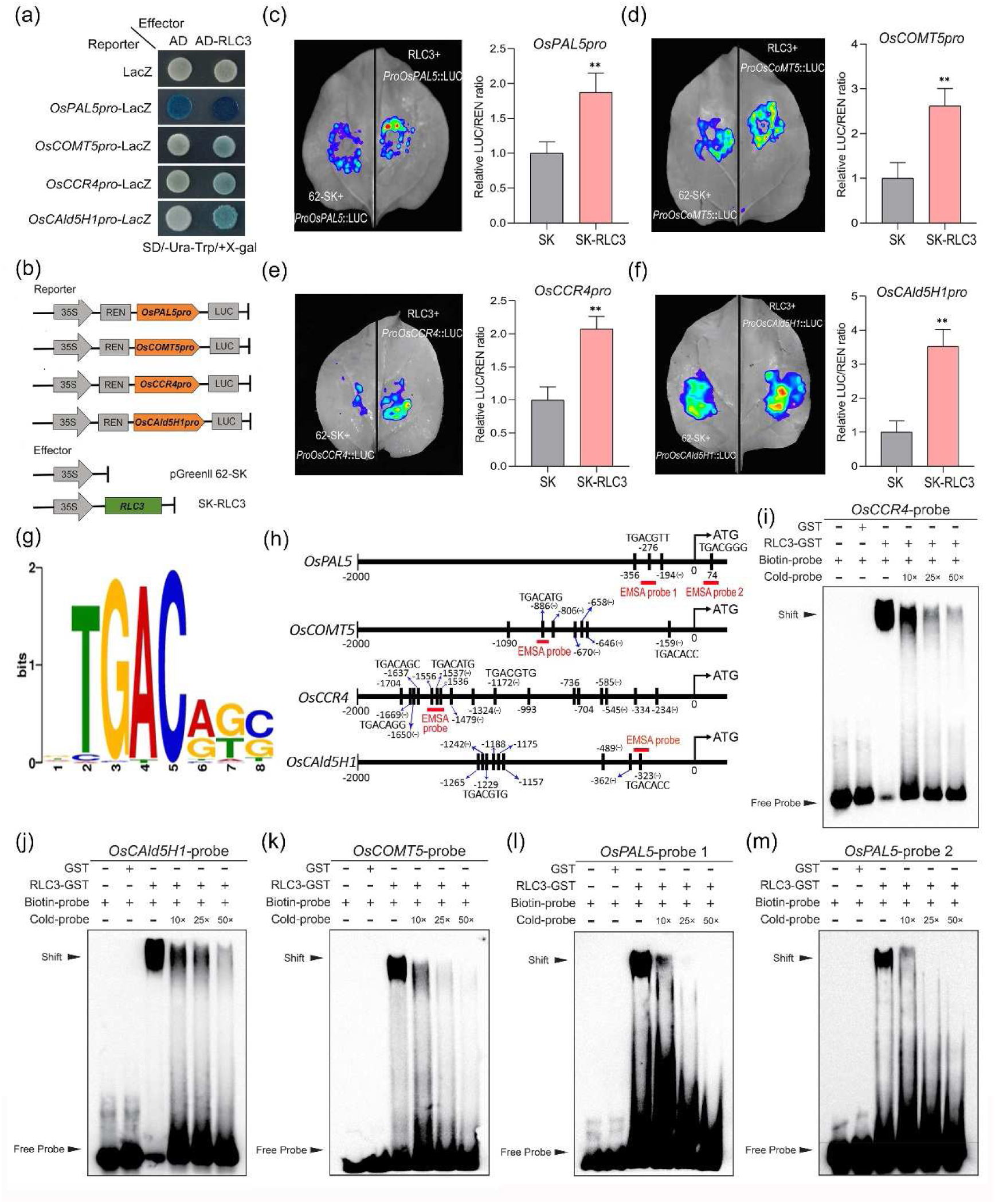
RLC3 directly bind to the promoters of lignin biosynthetic genes (*OsPAL5*, *OsCOMT5*, *OsCCR4*, and *OsCAld5H1*) and activate their transcription. (a) Yeast one-hybrid (Y1H) assay confirmed the interaction between RLC3 and the promoter of *OsPAL5*, *OsCOMT5*, *OsCCR4*, and *OsCAld5H1*. The effectors were AD and AD-RLC3, while the reporters were LacZ and promoter-LacZ. The control combinations were AD + LacZ, AD-RLC3 + LacZ, and AD + promoter-LacZ. (b) Schematic diagram of the reporter and effector constructs used in the Dual-LUC assay. (c-f) Dual-LUC assays showing the effects of RLC3 on *OsPAL5* (c), *OsCOMT5* (d), *OsCCR4* (e), and *OsCAld5H1* (f) promoter activity. The Dual-Luciferase assay was employed to measure the relative LUC/REN ratio, with higher values indicating stronger RLC3-mediated activation of the respective promoters. The LUC/REN ratio of the empty vector (SK) + promoter-LUC combination was normalized to 1. Data represent means ± SD from three independent replicates. Asterisks denote significant differences (Student’s t-test, \*\**P* < 0.01). (g) The consensus DNA-binding motif of TALE family transcription factors (BELL and KNOX). (h) Schematic representation of the DNA-binding motif in the promoter regions of *OsPAL5*, *OsCOMT5*, *OsCCR4*, and *OsCAld5H1*. The black horizontal line indicates the candidate gene promoter, while the black short vertical lines and corresponding numbers denote the positions of the binding motifs. The positions for EMSA probes are shown with red short horizontal line. (i-m) EMSA analysis of RLC3 binding to the promoters of *OsCCR4* (i), *OsCOMT5* (j), *OsCAld5H1* (k), *OsPAL5* (l), and the coding region of *OsPAL5* (m). The probes for each assay were labeled with biotin and incubated with recombinant RLC3-GST protein. Negative controls (no protein added or GST protein added) and competition assays with unlabeled probe (10×, 25× and 50× excess) were shown.

As a member of the TALE superfamily transcription factors, BELL and KNOX proteins share highly conserved binding motifs (TGAC) characteristic of this family (Fig. 9g). Motif scanning analysis in the JASPAR database revealed the presence of several TALE-specific binding motifs (TGAC) in the promoters of *OsPAL5* (>3), *OsCOMT5* (>7), *OsCCR4* (>17), and *OsCAld5H1* (>9) (Fig. 9h), suggesting these genes are probably direct transcriptional targets of RLC3/OsBLH4. Based on these findings, we designed electrophoretic mobility shift assays (EMSA) targeting the promoters of *OsPAL5*, *OsCOMT5*, *OsCCR4*, and *OsCAld5H1* (Fig. 9h). Our systematic investigation of these four genes began with *OsCCR4* due to its abundant TALE-specific binding motifs. The EMSA probe was designed to cover the 1521∼1566 nt region upstream of the start codon, encompassing three TGAC motifs (Fig. 9h), including the sequence TGACATG predicted by the JASPAR database to exhibit a very high binding affinity (relative score: 0.97). EMSA results revealed that RLC3 protein strongly binds to the *OsCCR4* promoter biotin-labeled probe, and this binding can be competitively inhibited by unlabeled cold probe (Fig. 9i). Similarly, EMSA experiments demonstrated that RLC3 protein strongly binds to biotin-labeled probes in the *OsCAld5H1* and *OsCOMT5* promoters, and can be competitively inhibited by cold probes (Fig. 9j-k). *OsPAL5* contains high-confidence TGAC motifs in both the promoter region and the downstream region of the start codon (74 nt, Fig. 9h). Therefore, we designed two biotin-labeled probes for EMSA validation. The results showed that RLC3 protein strongly binds to both probes, indicating direct binding on both the promoter and coding sequences of *OsPAL5* (Fig. 9l-m). Collectively, the EMSA results demonstrate that RLC3 specifically binds to the TGAC motifs in these DNA probes in vitro, thereby confirming physical interactions between RLC3 and the promoters of *OsPAL5*, *OsCOMT5*, *OsCCR4*, and *OsCAld5H1*. Taken together, these results support the regulatory role of RLC3 in activating *OsPAL5*, *OsCOMT5*, *OsCCR4*, and *OsCAld5H1* gene expressions, leading to enhanced lignin biosynthesis in rice.

## Discussion

The rice BELL family has 14 members (Mukherjee *et al*., 2009), with functionally characterized members including SH5/RI (Yoon *et al*., 2014; Ikeda *et al*., 2019), qSH1/RIL1 (Konishi *et al*., 2006; Zhang *et al*., 2009; Ikeda *et al*., 2019), OsBLH6 (Hirano *et al*., 2013), OsBIHD1 (Liu *et al*., 2017), and OsBLH4/CVD1/ROSES1 (Jing *et al*., 2017; Gong *et al*., 2018; Cao *et al*., 2024). These members regulate diverse developmental processes. Specifically, BELL-KNOX complexes control grain shattering through distinct mechanisms: the qSH1/SH5-OSH15 module represses lignin biosynthesis by suppressing *CAD2* (Yoon *et al*., 2017), while the qSH1-OSH71 complex activates *OsXTH12*, a xyloglucanase involved in hemicellulose formation (Wu *et al*., 2025). OsBLH6 also regulates lignin metabolism and secondary cell wall formation (Hirano *et al*., 2013). For *OsBLH4*, mutant analyses reveal pleiotropic roles: the *cvd1* mutant shows leaf rolling and affects plant height, heading date, and grain yield (Jing *et al*., 2017), while the *roses1* mutant exhibits reduced organ size, rolled leaves, elevated water loss, and accelerated senescence (Gong *et al*., 2018). CVD1/OsBLH4 controls plant height, grain number, and heading date by modulating *OsGA2ox1* promoter activity (Cao *et al*., 2024). A similar mechanism occurs in potato, where StBEL5-POTH1 heterodimers or individual proteins inhibit *GA20ox1* promoter activity (Chen *et al*., 2004). Despite these insights, the molecular basis of leaf rolling in *OsBLH4* remains unresolved.

In this study, we reveal that the BELL-type transcription factor RLC3/OsBLH4 interacts with KNOX transcription factors OSH1/45/71 to regulate lignin biosynthesis and secondary cell wall formation, thereby modulating leaf morphology and drought tolerance. We propose a working model in which the KNOX-BELL-lignin regulatory module mediates leaf rolling and drought resilience by initiating a multi-tiered regulatory cascade through two core mechanisms: direct activation of lignin biosynthetic genes and physical interaction with KNOX transcription factors (Fig. 10). RLC3 directly activates lignin biosynthetic genes (*OsPAL5*, *OsCOMT5*, etc.), driving secondary cell wall deposition. This process is further amplified by RLC3’s physical interaction with KNOX transcription factors (OSH1, OSH45, OSH71), forming a KNOX-BELL heterocomplex that synergistically enhances lignin content and cell wall rigidity. The enhanced lignin synthesis fortifies midrib structural integrity, ensuring efficient water transport through the vascular system. Disruption of this pathway compromises midrib function, leading to hydraulic imbalance and subsequent collapse of bulliform cells (BCs) due to impaired water retention. Ultimately, the cascading reinforcement of lignin deposition → cell wall rigidity → midrib integrity → water transport efficiency → BCs stability underpins RLC3’s critical role in optimizing leaf morphology and drought resilience by sustaining cellular turgor and photosynthetic activity under water deficit conditions.

**Fig. 10.**
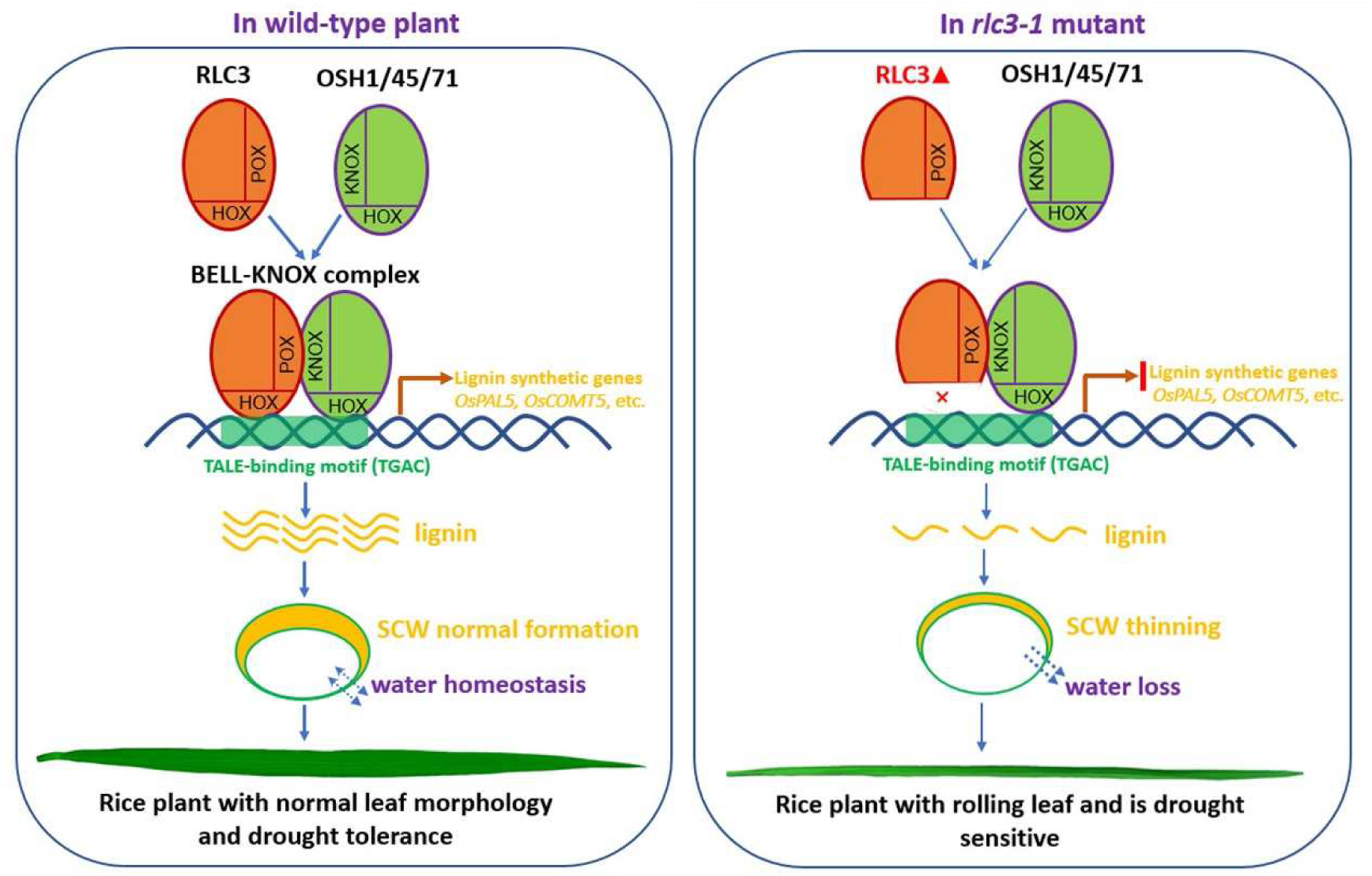
Working model of the KNOX-BELL-lignin regulatory module mediating leaf rolling and drought tolerance. (Left) In wild-type plants, RLC3 interacts with KNOX proteins OSH1/45/71 via its POX domain to form a BELL-KNOX complex. This complex binds to the TALE-binding motif (TGAC) in the promoters of lignin synthesis genes (*OsPAL5*, *OsCOMT5*, *OsCCR4*, and *OsCAld5H1*) through their homeobox (HOX) domains, promoting lignin synthesis and normal secondary cell wall (SCW) development in leaf midveins. This maintains normal leaf structure and water transport homeostasis, contributing to drought tolerance. (Right) In *rlc3-1* mutants, the mutation leads to the loss of the RLC3 protein’s HOX domain, which prevents binding to the TALE motif, blocking lignin gene expression and reducing lignin content. This results in SCW thinning, abnormal midvein structure, accelerated leaf water loss, bulliform cells dehydration, leaf rolling, and increased drought sensitivity.

### RLC3 mediates hydraulic-driven leaf rolling via coordinated BCs turgor and midrib integrity

BCs, specialized epidermal cells located between adjacent vascular bundles in the adaxial epidermis, control rice leaf rolling through turgor-dependent morphological changes (Xu *et al*., 2014, 2018; Li *et al*., 2010; Mai *et al*., 2024). Here, we identify *RLC3* as a master regulator of leaf rolling through map-based cloning and allelic complementation. The *RLC3* mutants (*rlc3-1*, *rlc3-2*, *rlc3-ko#11*, *rlc3-ko#25*) exhibit consistent reductions in BCs area and relative depth—phenotypes directly correlated with adaxial rolling—mirroring the cytological signature of *oslc1* (Wu *et al*., 2025). Crucially, these BCs defects are secondary to structural impairments in the midrib and secondary veins, which manifest as intense toluidine blue staining (Fig. 4), indicative of altered cell wall composition and accelerated water efflux. Concomitantly, leaf water content is significantly reduced (Fig. 5), confirming a hydraulic failure as the primary driver. Notably, epidermal cell morphology remains intact, distinguishing *rlc3* from *cld1/srl1* mutants, where epidermal staining dominates (Li *et al*., 2017). Unlike *sm1* (Jiang *et al*., 2025), which displays a rigid, solidified midrib causing abaxial rolling, *rlc3* mutants feature disorganized vascular conduits without mechanical reinforcement. We therefore propose a unified mechanistic model: *RLC3* governs midrib vascular integrity → maintains hydraulic conductivity → sustains BCs turgor → prevents adaxial rolling. This pathway redefines BCs as passive responders to vascular-derived water status, rather than autonomous morphogenetic units.

### RLC3 regulates drought tolerance by maintaining midrib structural integrity and hydraulic conductivity

Leaf morphology and structure, including BCs shape, epidermal architecture, and vascular patterning, directly determine water transport and retention capacity (Fang *et al*., 2012; Li *et al*., 2017; Mai *et al*., 2024). Under normal conditions, leaf rolling is strongly correlated with turgor pressure, primarily generated by BCs due to their superior hydraulic conductivity (Kadioglu *et al*., 2007; Matschi *et al*., 2020). However, during water stress, BCs rapidly dehydrate and shrink, losing structural support and causing leaf rolling. This process is reversible upon rehydration (Kadioglu *et al*., 2007). In this study, RLC3 mutant lines exhibited smaller BCs and reduced relative depth, directly contributing to decreased leaf water content and increased water loss rates compared to wild-type (WT) plants. Unlike *CLD1/SRL1* mutants, which link epidermal defects to water deficits (Li *et al*., 2017), toluidine blue staining revealed no epidermal differences between *rlc3* mutants and WT. Instead, all *rlc3* mutant midribs showed distinct blue staining, indicating localized structural defects impairing vascular function and water transport (Mai *et al*., 2024). This contrasts with *OsACL5* overexpression phenotypes, where reduced vascular size leads to similar water transport deficiencies (Mai *et al*., 2024). As the central integrative structure, the midrib system facilitates nutrient/water transport, structural support, gas exchange, and stress mitigation (Jiang *et al*., 2025). Our data establish midrib structural impairment as the primary cause of reduced water content and elevated water loss in *rlc3* mutants. *RLC3* regulates water transport by maintaining midrib structural integrity through secondary cell wall (SCW) formation. This is evidenced by significantly reduced SCW thickness in *rlc3* mutants, primarily due to diminished lignin deposition (Fig. 6). The *RLC3* mutation specifically impairs SCW formation in midrib vascular bundles, compromising structural integrity and water transport efficiency.

Leaf water loss is a critical determinant of drought tolerance (Chen *et al*., 2021; Li *et al*., 2023). *RLC3* loss-of-function mutations induce anatomical defects, including aberrant BCs morphology and compromised midrib structure, directly contributing to higher water loss rates and reduced relative water content (Fig. 5). These impairments manifest as increased electrolyte leakage, elevated malondialdehyde (MDA) levels, decreased chlorophyll content, and reduced proline accumulation. These phenotypes indicate severe cell membrane damage, diminished photosynthetic stability, and impaired osmoregulatory capacity, collectively reducing drought tolerance. This underscores RLC3’s pivotal role in drought stress responses. We propose that RLC3 sustains drought tolerance through two primary mechanisms: (1) regulation of leaf water transport and (2) maintenance of leaf structural integrity—both essential for normal growth and development. This conclusion aligns with our earlier work on *CLD1/SRL1* mutants (Li *et al*., 2017), where BCs shrinkage due to accelerated water loss triggers drought response pathways, promoting chlorophyll accumulation and sustaining normal growth under stress.

### RLC3 activates lignin biosynthesis to ensures midrib hydraulic function and drought resilience

Lignin, a major plant cell wall component, critically determines mechanical strength and structural stability. Previous studies have linked cell wall composition—particularly lignin and cellulose content—to rice leaf rolling phenotypes (Luan *et al*., 2011; Fang *et al*., 2012; Yang *et al*., 2014; Li *et al*., 2017). To investigate this further, we conducted transcriptome sequencing on wild-type and *rlc3-1* mutant plants under normal conditions. The analysis revealed significant enrichment of differentially expressed genes (DEGs) in lignin biosynthesis pathways, suggesting *RLC3* plays a key regulatory role in lignin synthesis. Supporting this, lignin quantification, histochemical staining, and TEM observations confirmed reduced lignin accumulation and SCWs in *rlc3* mutant midrib vascular bundles compared to wild-type plants (Fig. 6). These findings align with previous *RL14* studies (Fang *et al*., 2012), where impaired lignin and cellulose content disrupted cell wall integrity, compromised water transport, and induced leaf rolling. Given the midrib structural defects observed in *rlc3* mutants, we propose that defective lignin biosynthesis underlies the leaf water deficit and rolling phenotype by weakening vascular tissue integrity.

Extensive research has demonstrated that lignin accumulation significantly enhances water deficit tolerance across diverse plant species, including rice (Bang *et al*., 2019, 2022; Jung *et al*., 2022), maize (Hu *et al*., 2009), wheat (Luo *et al*., 2023), and apple (Hou *et al*., 2022; Jing et al., 2024). Consistent with these findings, our study revealed that all *rlc3* mutant lines exhibited markedly increased drought sensitivity and reduced lignin content compared to wild-type plants, suggesting *RLC3* deficiency decreases lignin content in leaves, weakening cell wall structure and reducing water retention, which collectively account for the decreased drought tolerance observed in *rlc3* mutants. Further mechanistic analysis identified RLC3 as a transcription factor (OsBLH4) that directly binds to promoters of lignin biosynthesis genes (*OsPAL5*, *OsCOMT5*, *OsCCR4*, and *OsCAld5H1*). Importantly, loss-of-function mutations in *RLC3* resulted in depressed expression of these genes, leading to reduced lignin accumulation and subsequent cell wall structural defects. Collectively, these findings establish RLC3 as a positive regulator of lignin biosynthesis and SCW formation. The observed drought hypersensitivity in *rlc3* mutants underscores the critical role of RLC3-mediated lignin deposition in maintaining SCW formation, cellular water balance, and stress resilience.

### RLC3-OSH1/45/71 heterodimers coordinate lignin biosynthesis and vascular integrity to govern leaf morphogenesis and drought tolerance

The evolutionarily conserved BELL-KNOX heterodimer system governs fundamental developmental processes across plant species, including cell proliferation, organ morphogenesis, and hormone signaling (Müller *et al*., 2001; Chen *et al*., 2004; Kanrar *et al*., 2006; Yoon *et al*., 2017; Wu *et al*., 2025). Our study extends this paradigm by experimentally validating specific interactions between the BELL protein RLC3/OsBLH4 and the KNOX proteins OSH1, OSH45, and OSH71 in rice, with the POX domain of RLC3 identified as the critical interface. This finding reinforces the functional conservation of the BELL-KNOX partnership while pinpointing its role in leaf development.

Genetic analysis of KNOX mutants revealed distinct yet informative leaf phenotypes that implicate RLC3 interactions. Both *osh1* and *osh45* knockout lines exhibited mild leaf curling, partially phenocopying the leaf rolling observed in *rlc3* mutants. This phenotypic convergence strongly suggests a shared functional module involving RLC3, OSH1, and OSH45 in regulating leaf flatness. In contrast, *osh71* mutants displayed a unique, flattened, and shortened leaf morphology without curling, distinct from the *rlc3* mutant, underscoring OSH71’s critical role in leaf development. This divergence suggests that OSH71 operates within a more complex regulatory network, potentially involving crosstalk with additional pathways, consistent with its pleiotropic roles exemplified by its partnership with qSH1 in activating *OsXTH12* to control seed shattering (Chen *et al*., 2023).

Ectopic expression of various KNOX genes (e.g., *OSH1*, *OSH6*, *OSH15*, *OSH43*, *OSH71*) causes a spectrum of developmental abnormalities in rice, from severe (green organs with ectopic buds) to moderate (absent normal leaves) and mild (normal leaves/sheaths but lacking ligules with diffuse boundaries) (Sentoku *et al*., 2000). This demonstrates KNOX genes’ essential role in rice leaf morphogenesis and suggests their dosage-dependent function in coordinating development.

Building on the established role of the OsbHLH002-OSH1-OsNAC31 module in regulating cellulose deposition and SCW thickening in leaf sheaths (Chen *et al*., 2023), our work provides a crucial mechanistic expansion. We demonstrate that OSH1, along with OSH45 and OSH71, is also indispensable for lignin biosynthesis. Mutations in these KNOX genes resulted in reduced lignin content and attenuated SCW thickness in midrib vascular bundles. Collectively, our data establish that RLC3, in concert with OSH1/45/71 through the BELL-KNOX module, coregulates a suite of lignin biosynthesis genes (e.g., *OsPAL5*, *OsCOMT5*, *OsCCR4*, *OsCAld5H1*). This coordinated regulation is essential for proper midrib SCW assembly.

Therefore, we propose a cohesive mechanistic framework: the RLC3-KNOX heterodimer functions as a master regulator of SCW formation, synchronizing lignin biosynthesis with vascular integrity. This mechanism directly determines midrib vascular integrity, which in turn governs hydraulic conductivity and leaf water status—ultimately shaping leaf morphology and conferring drought resilience. This model aligns with broader observations of BELL-KNOX-mediated repression of lignin genes in other contexts (e.g., OSH15-qSH1 suppressing *CAD2* in shattering) but uniquely positions the RLC3-centered module as a critical integrator linking developmental patterning with abiotic stress adaptation. Our findings elucidate that RLC3’s influence extends beyond mere morphological control, embedding drought tolerance as a core physiological outcome of its SCW regulatory network.

## Acknowledgements

This work was supported by the National Natural Science Foundation of China (grant nos. 32070197) and the Key Research and Development Program of Shaanxi Province (2023-YBNY-089). We extend our gratitude to Prof. Dr. Juaner Dong (College of Life Sciences, Northwest A&F University) for generously providing protein expression plasmids essential for this study. Additionally, we acknowledge the technical support provided by Xiyan Chen, Ningjuan Fan and Zhongping Lei (Teaching and Research Core Facility at College of Life Sciences, Northwest A&F University) in microscopic observation, fluorescence imaging, and protein purification experiments.

## Author contributions

WL conceived the study and designed the experiments. LQ and ZZ performed most of the experiments and analyzed the data. QL, KD, JL, ML, YC, ZL, and LZ performed some experiments. HL and KC contributed to experimental design. LQ wrote the manuscript, and WL revised it. LQ and ZZ contributed equally to this work.

## Supporting Information

**Fig. S1.** Genetic mapping of the *rlc3-2* mutant and analysis of mutation sites in *rlc3-1* and *rlc3-2*.

**Fig. S2.** Sanger sequencing chromatograms of mutation sites in parental lines and F_1_ progeny from reciprocal crosses among *rl3-1*, *rlc3-2*, and *rl3-ko#25* mutants.

**Fig. S3.** Phenotypic characterization and leaf curling quantification in parental lines and F_1_ progeny from reciprocal crosses among *rl3-1*, *rlc3-2*, and *rl3-ko#25* mutants.

**Fig. S4.** Comparative analysis of protein domain architecture and predicted 3D structures of RLC3 and its alternatively spliced isoforms RLC3-v2 and RLC3-v3.

**Fig. S5.** Multiple sequence alignment and phylogenetic analysis of RLC3 and its homologs across plant species.

**Fig. S6.** Subcellular localization and tissue-specific expression profiles of RLC3-v2 and RLC3-v3 protein isoforms.

**Fig. S7.** Paraffin-embedded cross-sections of leaves from wild-type and *rlc3* mutant lines.

**Fig. S8.** Comparison of drought tolerance during grain-filling stage between wild-type and *rlc3* mutant lines.

**Fig. S9.** GO enrichment analysis of differentially expressed genes (DEGs) identified in the transcriptome sequencing of wild-type versus *rlc3-1* mutant.

**Fig. S10.** KEGG pathway analysis of DEGs involved in phenylpropanoid biosynthesis in wild-type versus *rlc3-1* mutant.

**Fig. S11.** KEGG pathway analysis of DEGs involved in flavonoid biosynthesis in wild-type versus *rlc3-1* mutant.

**Fig. S12.** KEGG pathway analysis of DEGs involved in phenylalanine, tyrosine, and tryptophan biosynthesis in wild-type versus *rlc3-1* mutant.

**Fig. S13.** Prediction of RLC3-interacting proteins using the PRIN database.

**Fig. S14.** Subcellular co-localization analysis of RLC3 with OSH1, OSH45, and OSH71 proteins.

**Fig. S15.** Comparison of flag leaf length, width, and plant height at maturity between wild-type and *OSH1*, *OSH45*, and *OSH71* knockout mutants.

**Fig. S16.** Expression analysis of lignin biosynthesis genes (*OsPAL5*, *OsCOMT5*, *OsCCR4*, and *OsCAld5H1*) in wild-type and *rlc3* mutant lines.

**Table S1.** DNA primers used in this study.

**Table S2.** Statistical analysis of agronomic traits in wild-type and *rlc3-1*, *rlc3-2* mutants.

**Table S3.** Comprehensive list of differentially expressed genes (DEGs) identified by transcriptome sequencing between wild-type and *rlc3-1* mutant.

**Table S4.** Lignin-related differentially expressed genes (DEGs) identified by transcriptome sequencing in comparisons between wild-type and *rlc3-1* mutant.

