## Supplemental Figures S1-S16 for "The BELL-type homeobox transcription factor RLC3/OsBLH4 controls leaf rolling and drought tolerance via KNOX-BELL-lignin regulatory network in rice"

### Supporting Information

### Fig. S1~S16

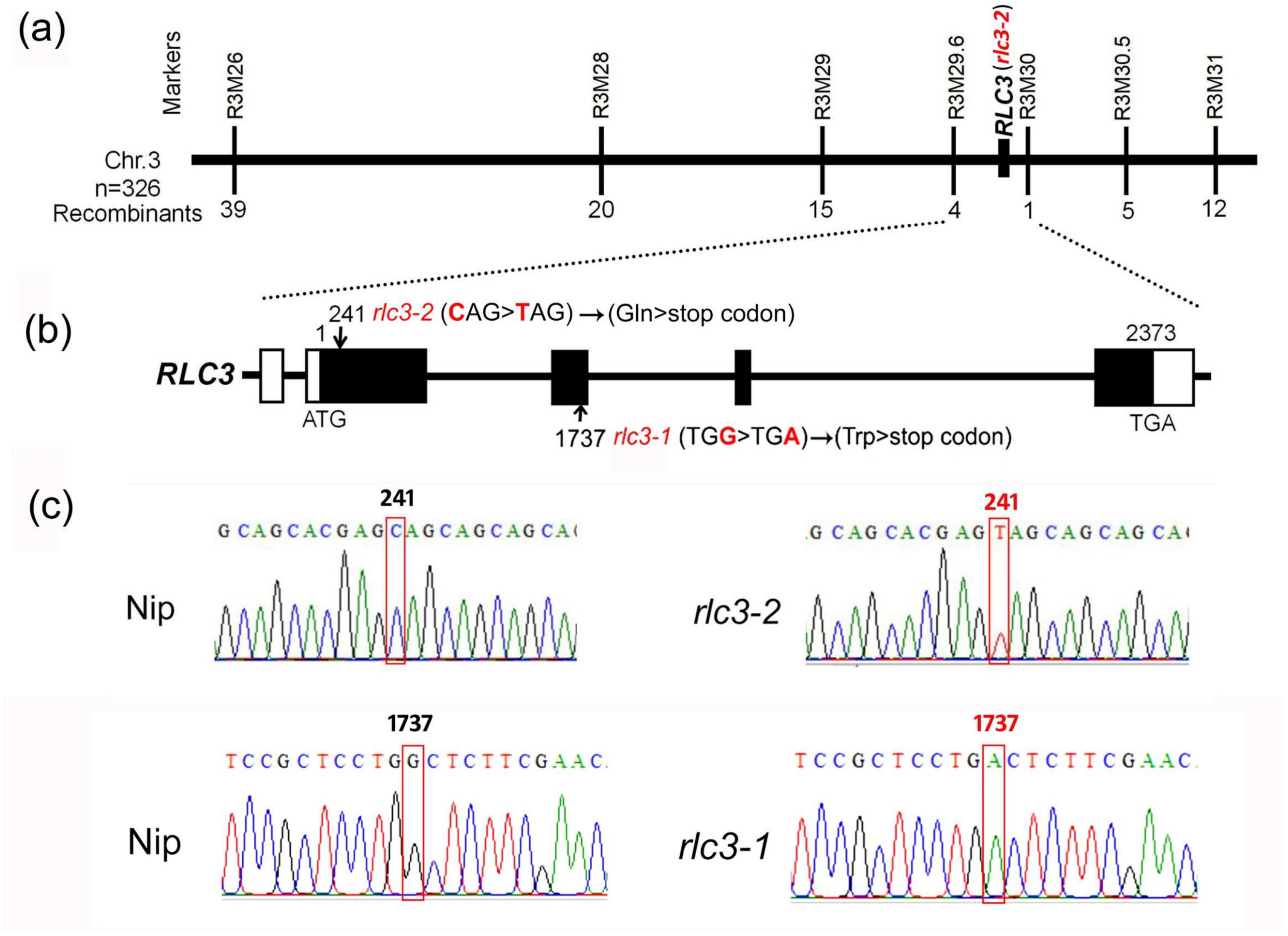


**Fig. S1.** Genetic mapping of the *rlc3-2* mutant and analysis of mutation sites in *rlc3-1* and *rlc3-2*. (a) *RLC3* gene localization linkage map. Markers' names are above the vertical lines and the number of recombinants is displayed under the vertical lines. (b) Gene structure and mutant sites of candidate gene *LOC_Os03g52239* (*RLC3*). Black boxes represent exons, horizontal lines represent introns, and white boxes represent UTR regions. (c) Peak sequencing of *RLC3* gene mutation sites in wild-type, *rlc3-1,* and *rlc3-2* mutants.

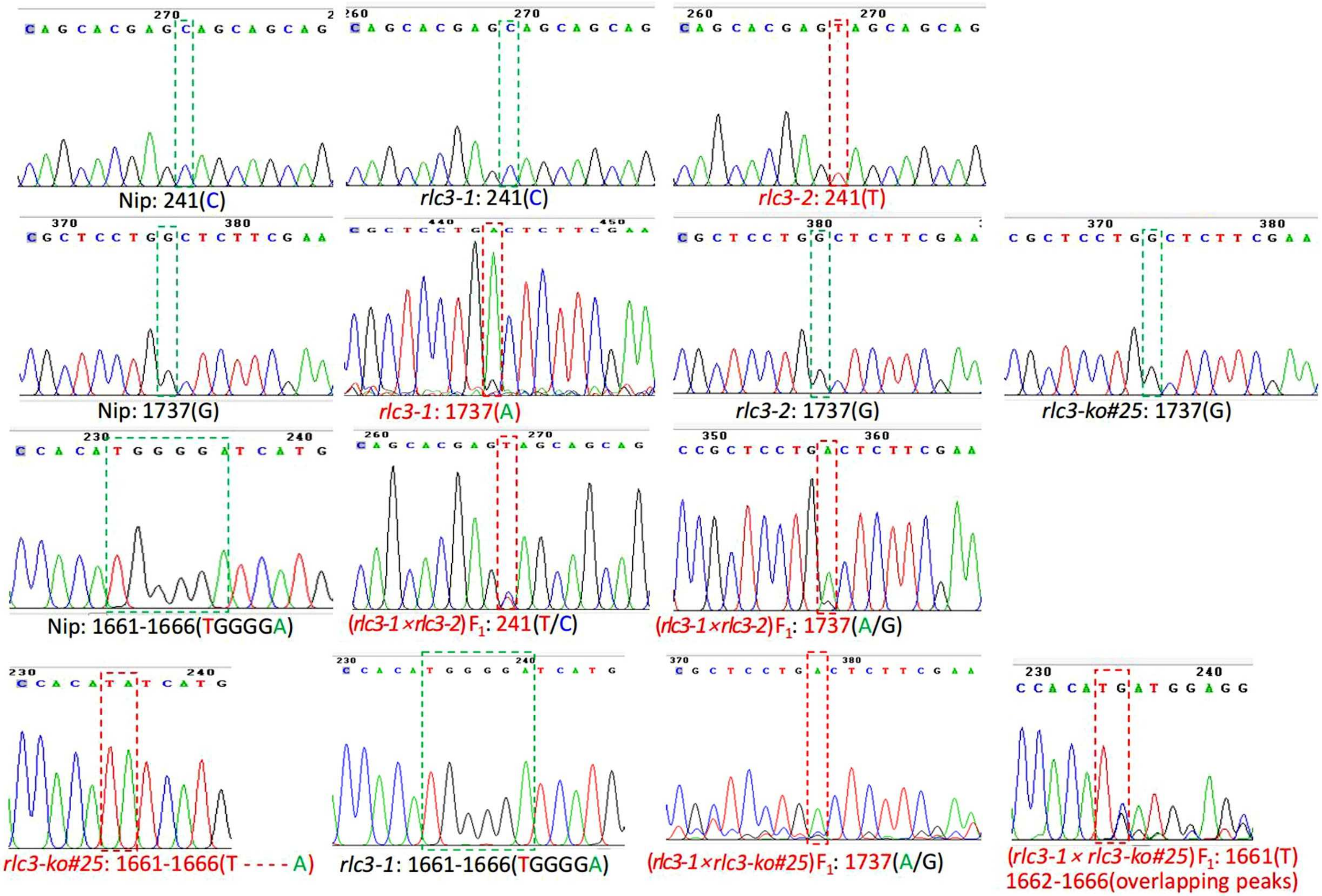


**Fig. S2.** Sanger sequencing chromatograms of mutation sites in parental lines and $F_1$ progeny from reciprocal crosses among *rl3-1*, *rlc3-2*, and *rl3-ko#25* mutants.

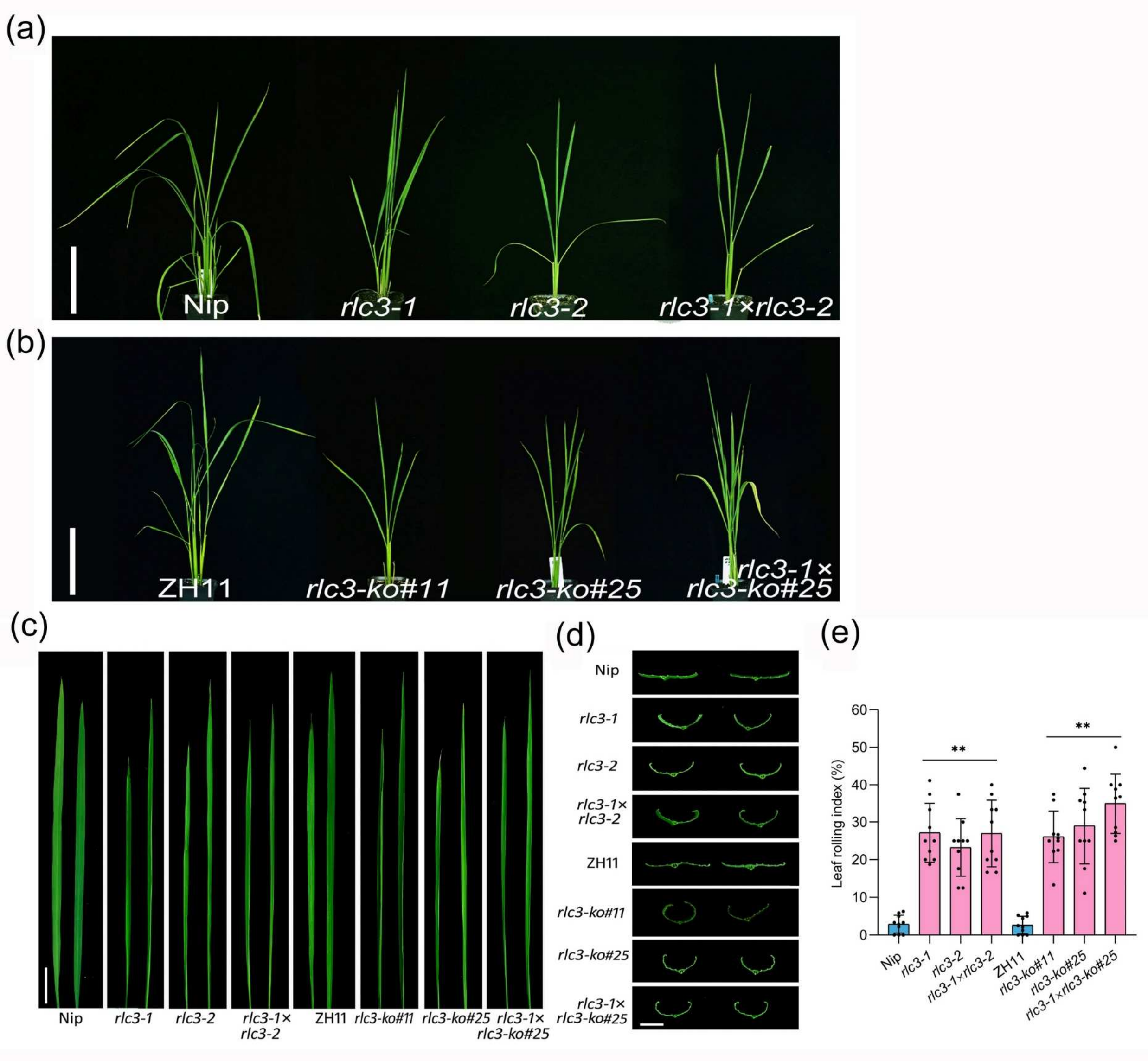


**Fig. S3.** Phenotypic characterization and leaf curling quantification in parental lines and $F_1$ progeny from reciprocal crosses among *rl3-1*, *rlc3-2*, and *rl3-ko#25* mutants. (a) The gross phenotype of the wild type Nip, mutants *rlc3-1* and *rlc3-2*, and the hybrid $F_1$ generation *rlc3-1*×*rlc3-2* plants. Bar=15 cm. (b) The gross phenotype of the wild type ZH11, knockout lines *rlc3-ko#11* and *rlc3-ko#25*, and the hybrid $F_1$ generation *rlc3-1*×*rlc3-ko#25* plants. Bar=15 cm. (c) The leaf phenotypes of wild-type, mutant and hybrid $F_1$ plants. Bar=3 cm. (d) The leaf blades crosssection of wild-type, mutants and hybrid $F_1$ plants. Bar=0.5 cm. (e) The leaf rolling index of wild-type, mutants and hybrid $F_1$ plants. n=10, Data are means ± SD, asterisks indicate significant differences according to Student's *t*-test (***P* < 0.01).

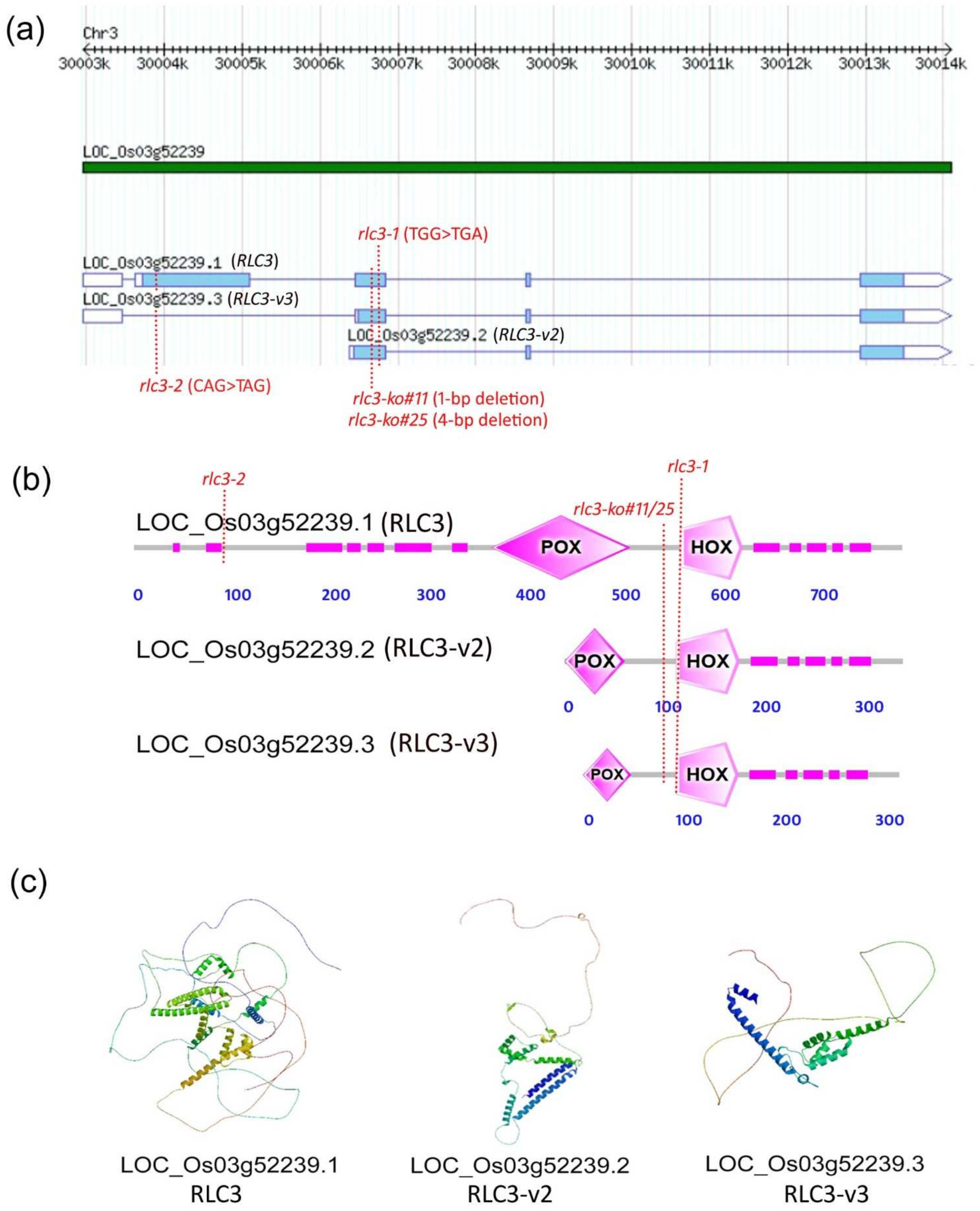


**Fig. S4.** Comparative analysis of protein domain architecture and predicted 3D structures of RLC3 and its alternatively spliced isoforms RLC3-v2 and RLC3-v3. (a) The gene structure of the three transcripts of *RLC3* gene.The white boxes or white arrows indicate the UTR regions, the blue boxes represent the exon regions, the horizontal lines signify the intron regions, and the scale at the top shows the physical distance of the gene. (b) *RLC3* gene three-transcript protein structure. (c) *RLC3* gene three-transcript protein threedimensional structure model.

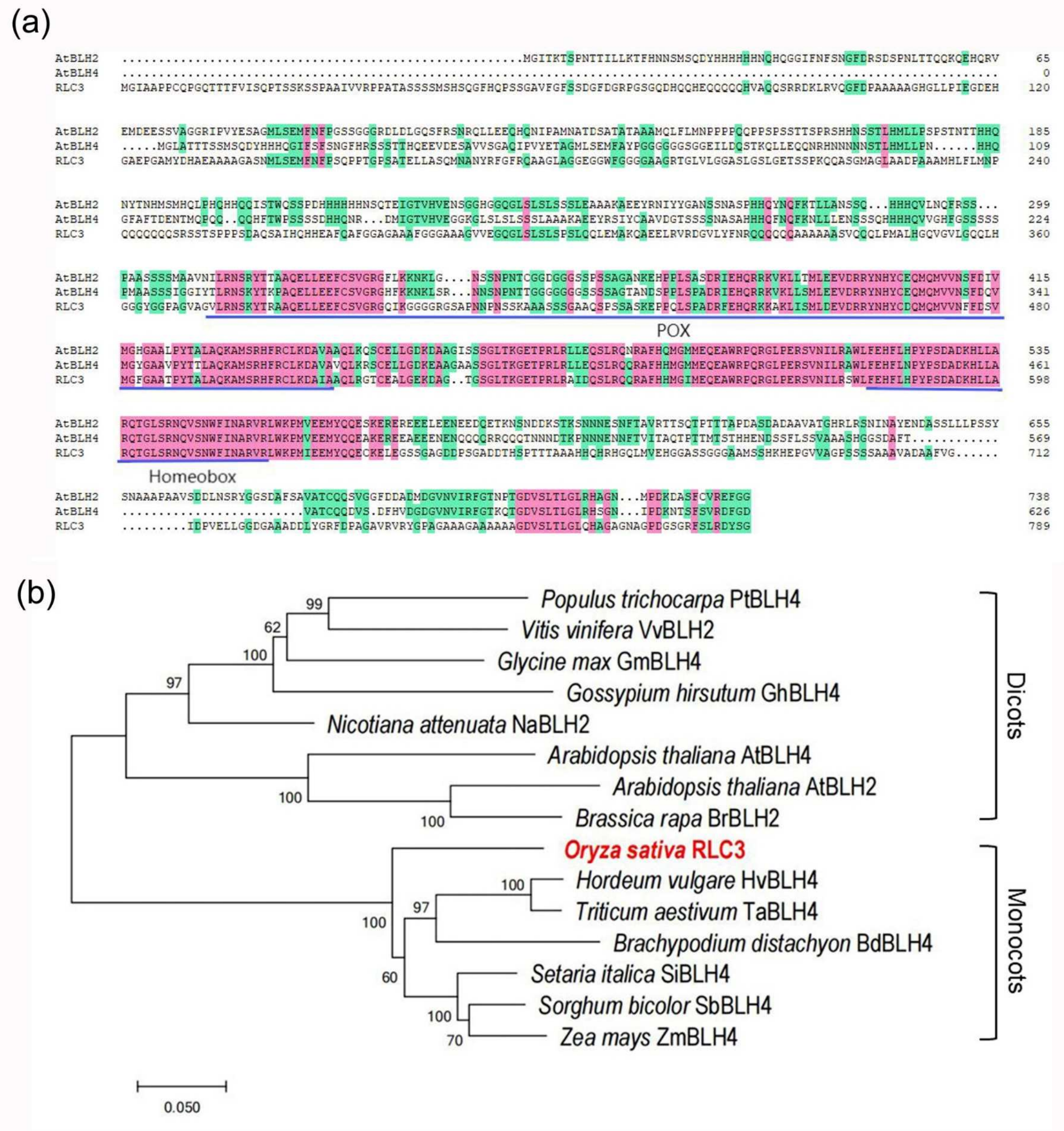


**Fig. S5.** Multiple sequence alignment and phylogenetic analysis of RLC3 and its homologs across plant species. (a) Sequence alignment analysis was performed using the full-length amino acid sequences of RLC3, as well as BLH2 and BLH4 from Arabidopsis thaliana, with POX and Homeobox being the conserved domains as shown in the figure. (b) The phylogenetic tree was constructed using RLC3 and homologous sequences from 13 species, with node values in the phylogenetic tree derived from 1,000 bootstrap replicates.

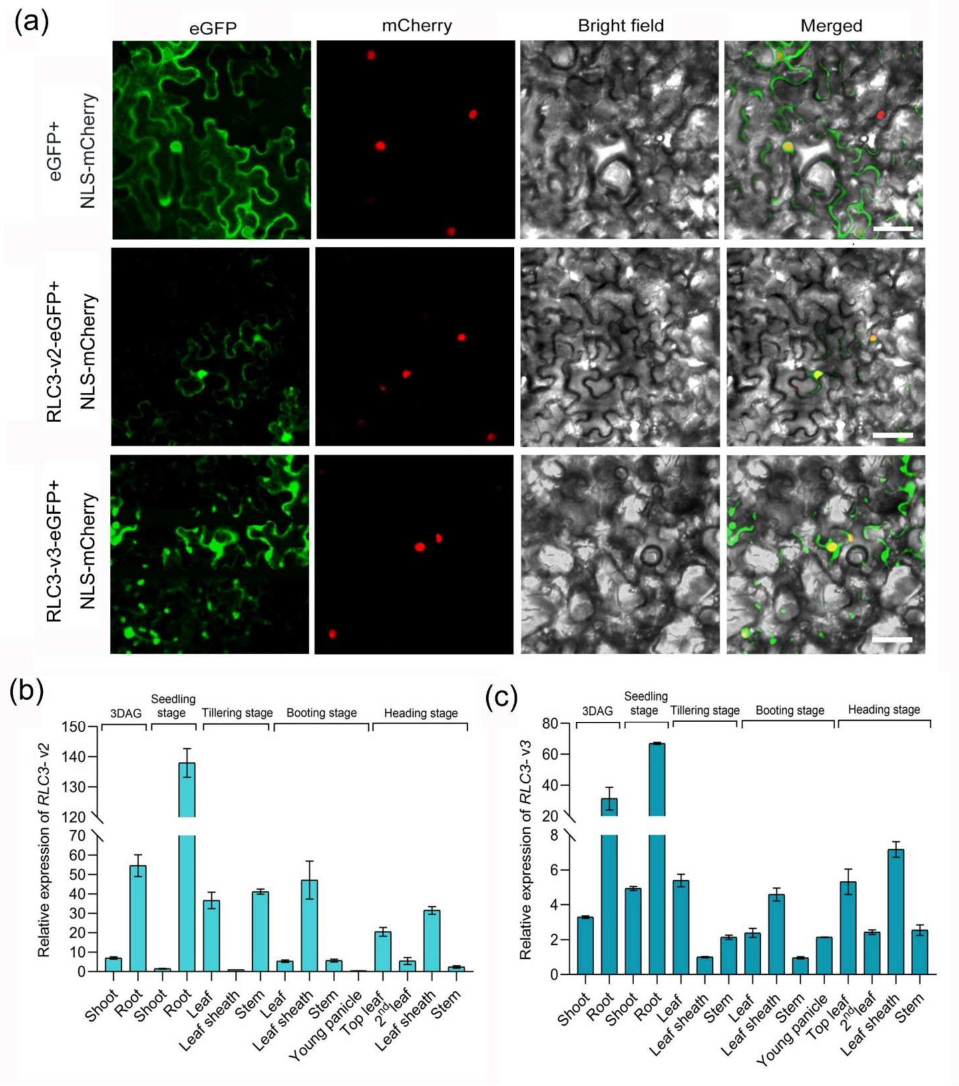


**Fig. S6.** Subcellular localization and tissue-specific expression profiles of RLC3-v2 and RLC3-v3 protein isoforms. (a) Subcellular localization of p35S-eGFP, RLC3-v2 (LOC_Os03g52239.2), and RLC3-v3 (LOC_Os03g52239.3) proteins. (b-c) The tissue expression profiles of RLC3-v2 (LOC_Os03g52239.2) and RLC3-v3 (LOC_Os03g52239.3). *Ubiquitin* (*Ubq*) as the internal control gene. The expression level in the leaf sheath at the tillering stage was set as 1. The relative expression levels of the gene were calculated using the $2^{-\Delta\Delta Ct}$ method. The experiment was conducted with three biological replicates and three technical replicates. Data are means ± SD.

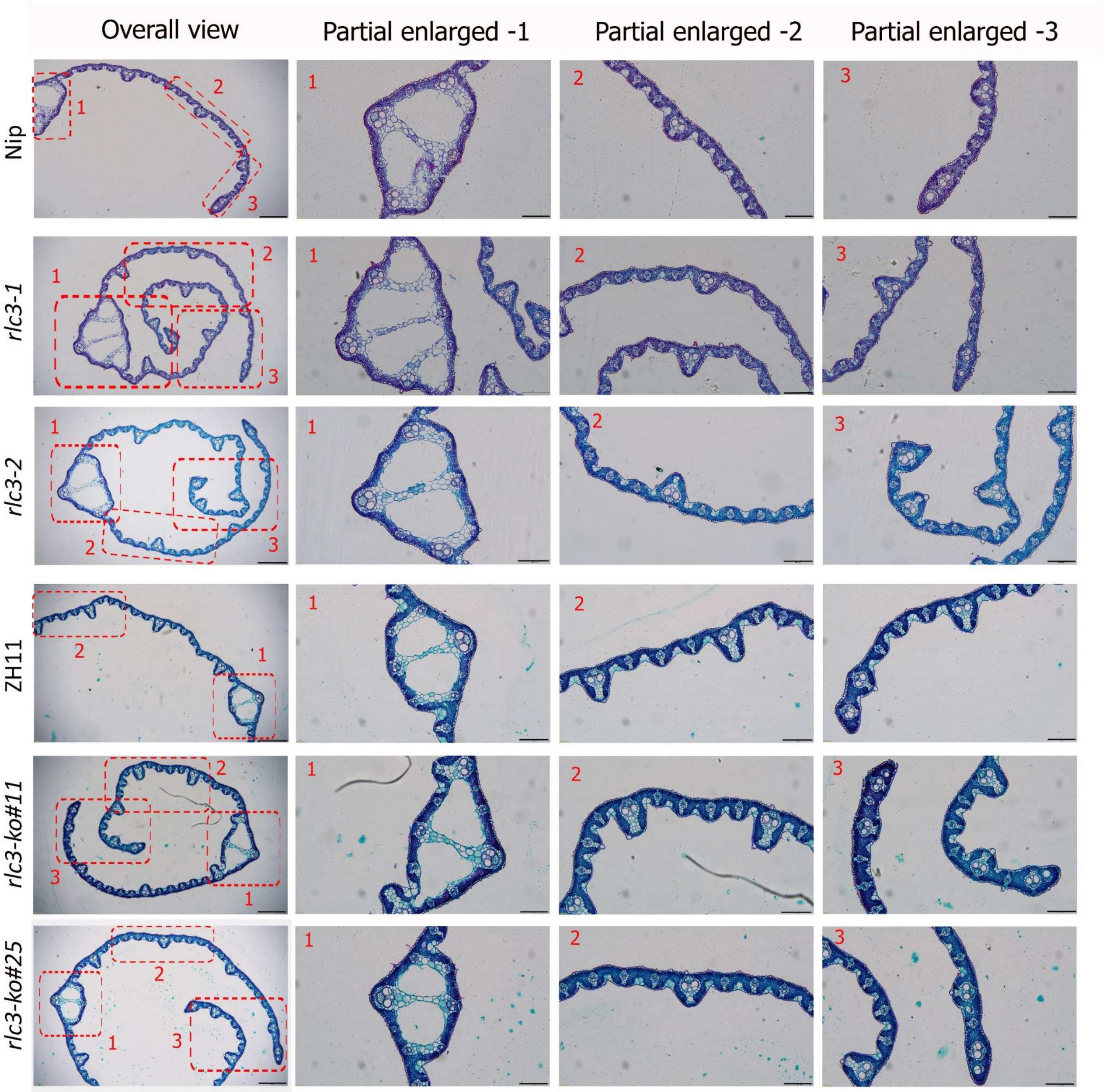


**Fig. S7.** Paraffin-embedded cross-sections of leaves from wild-type and *rlc3* mutant lines.

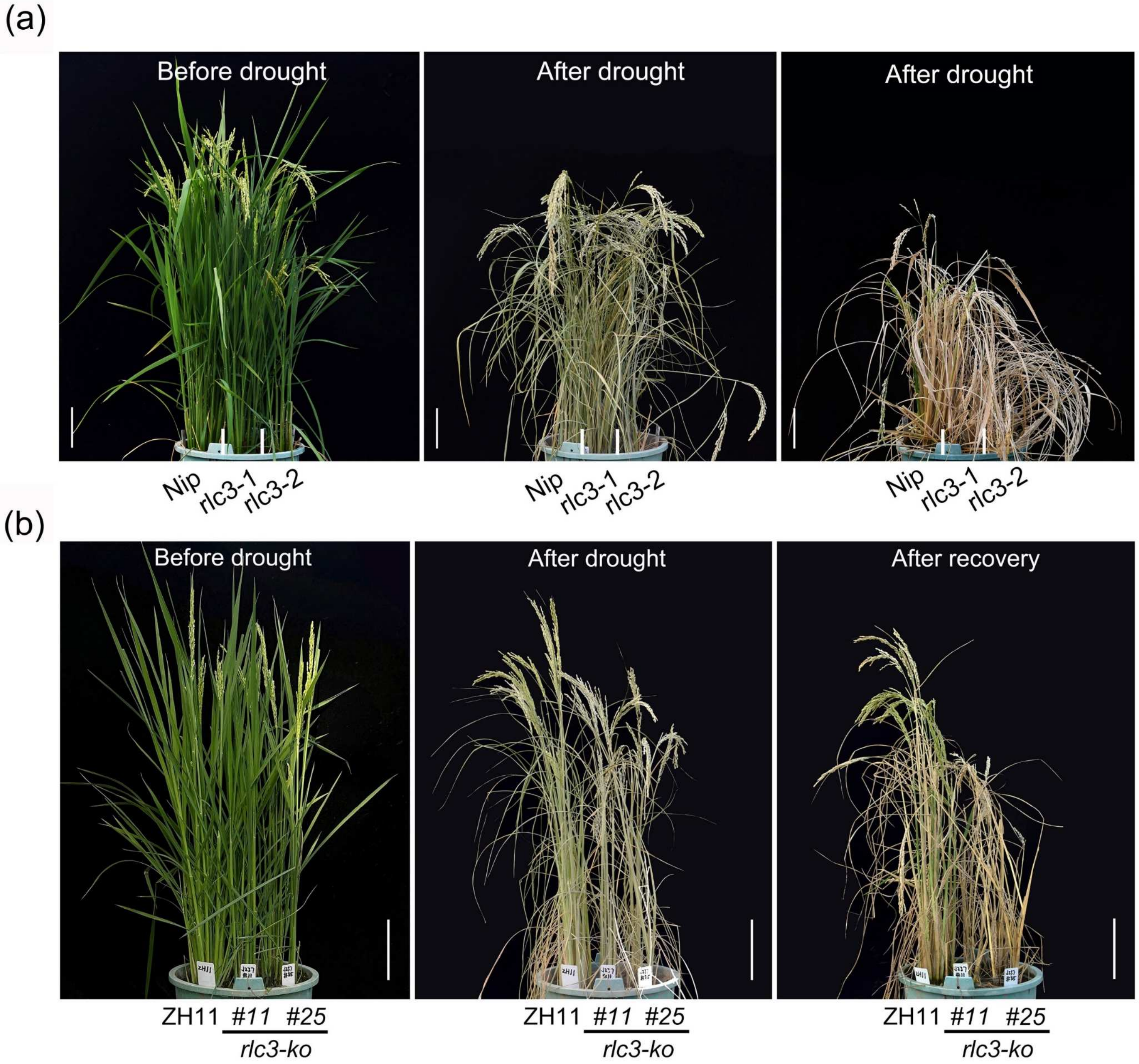


**Fig. S8.** Comparison of drought tolerance during grain-filling stage between wild-type and *rlc3* mutant lines. (a) Phenotypes of wild-type Nip and *rlc3-1* and *rlc3-2* mutants after drought stress treatment during grain filling stage. bars=15 cm. (b) Phenotypes of wild-type ZH11 and knockout lines *rlc3-ko#11* and *rlc3-ko#25* after drought stress treatment during grain filling stage. bars=20 cm.

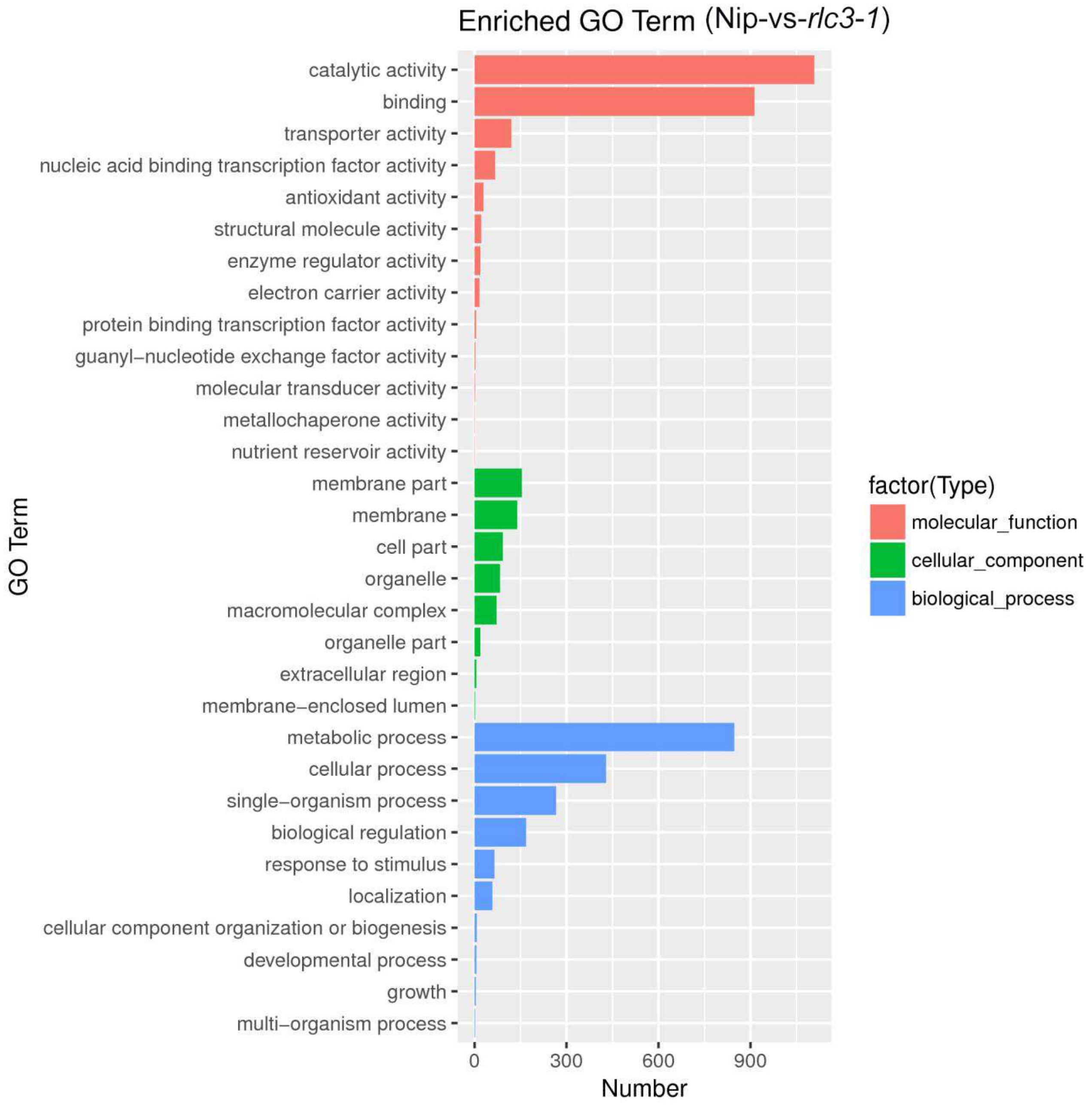


**Fig. S9.** GO enrichment analysis of differentially expressed genes (DEGs) identified in the transcriptome sequencing of wild-type versus *rlc3-1* mutant. The vertical axis represents enriched GO terms, the horizontal axis represents the number of differentially expressed genes. Different colored bar charts represent molecular functions, cellular components, and biological processes.

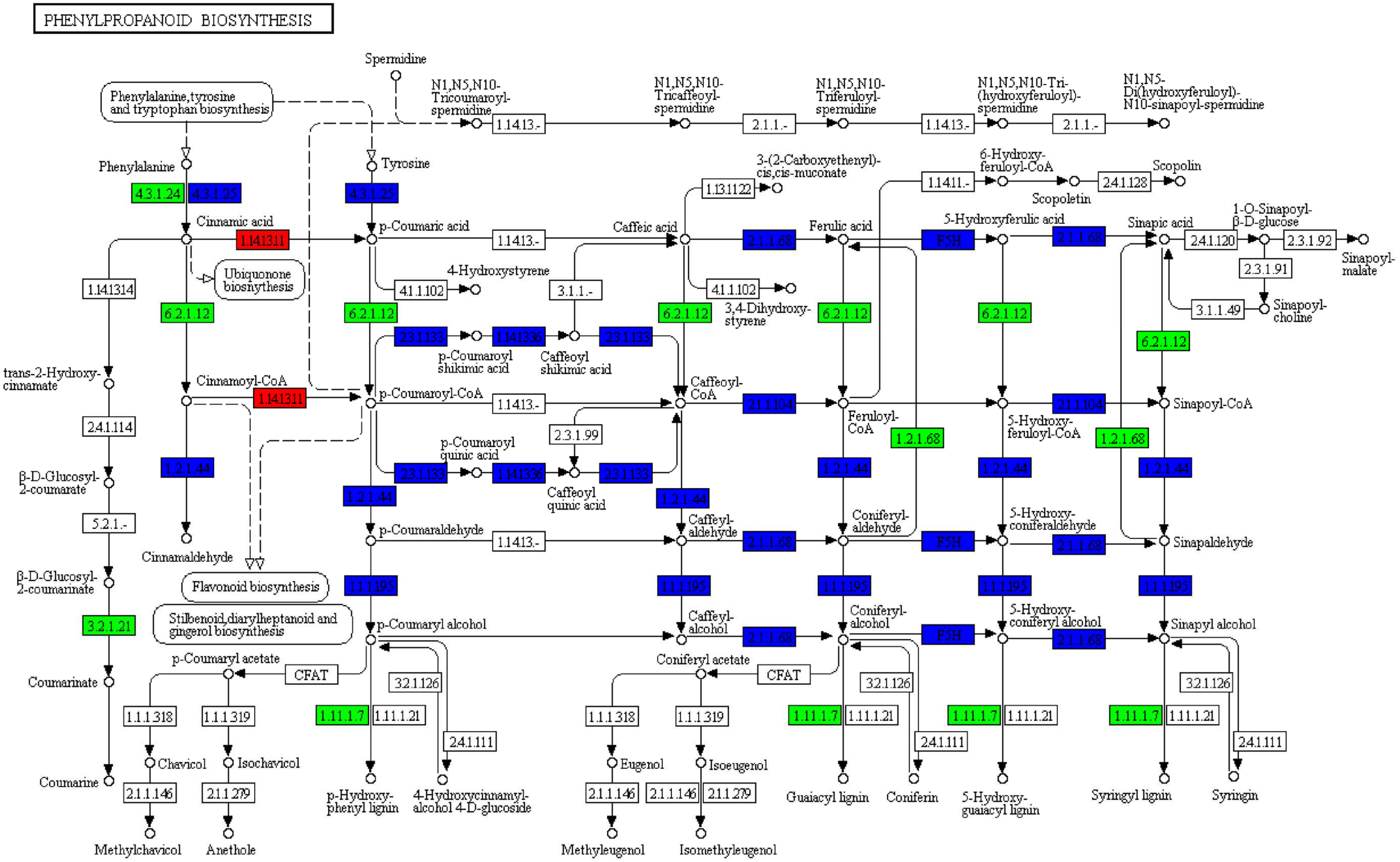


**Fig. S10.** KEGG pathway analysis of DEGs involved in phenylpropanoid biosynthesis in wild-type versus *rlc3-1* mutant.

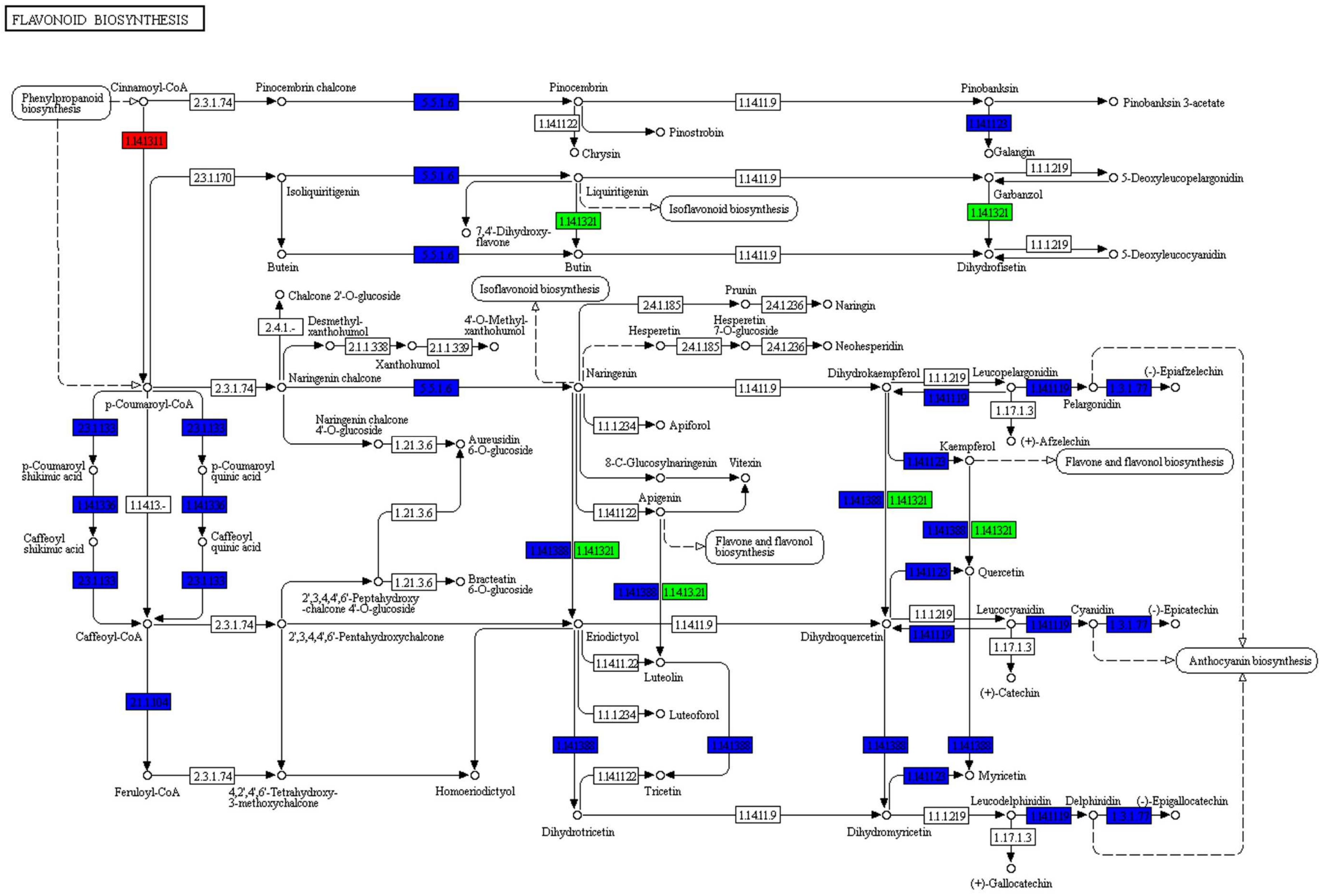


**Fig. S11.** KEGG pathway analysis of DEGs involved in flavonoid biosynthesis in wild-type versus *rlc3-1* mutant.

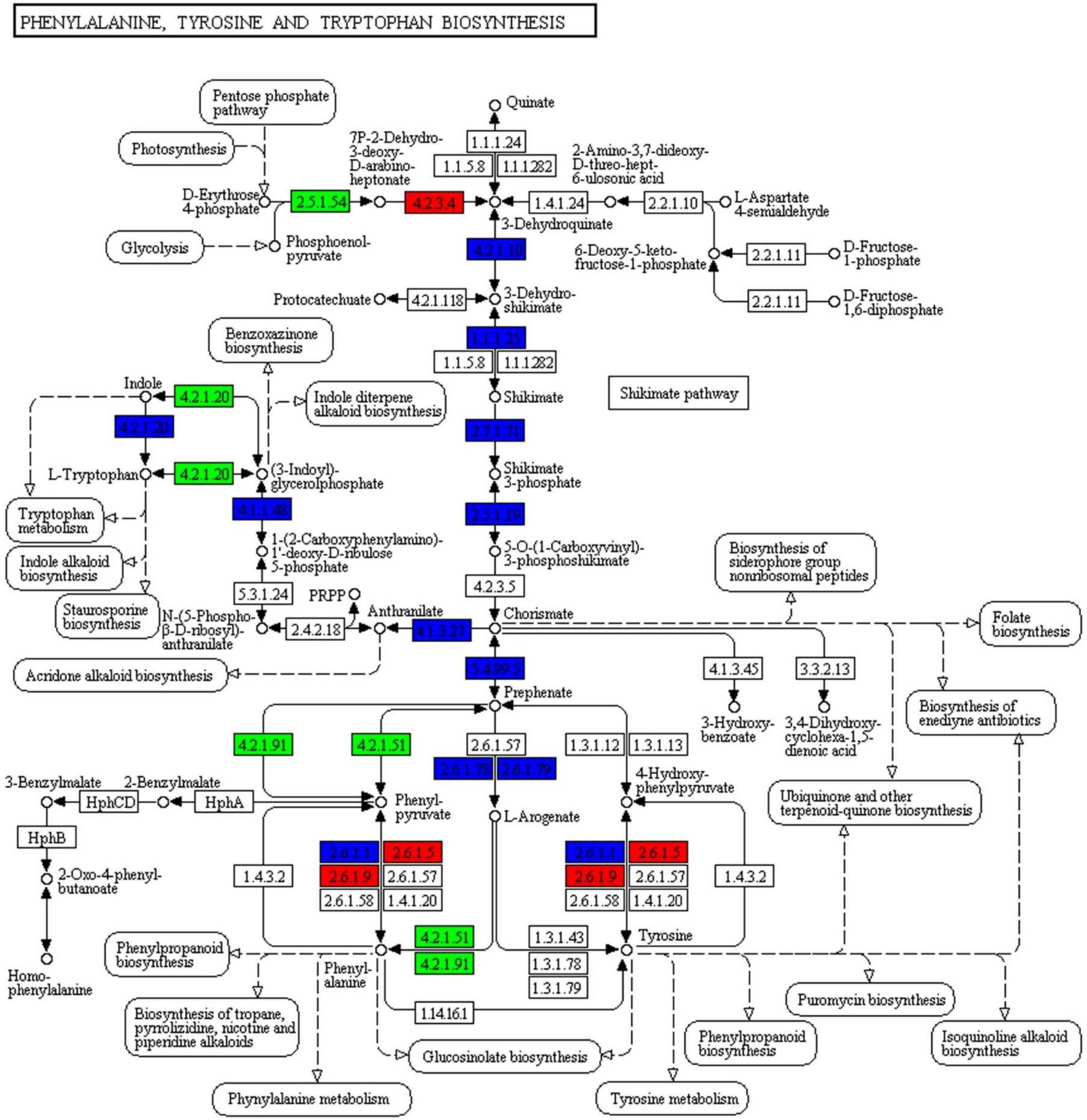


**Fig. S12.** KEGG pathway analysis of DEGs involved in phenylalanine, tyrosine, and tryptophan biosynthesis in wild-type versus *rlc3-1* mutant.

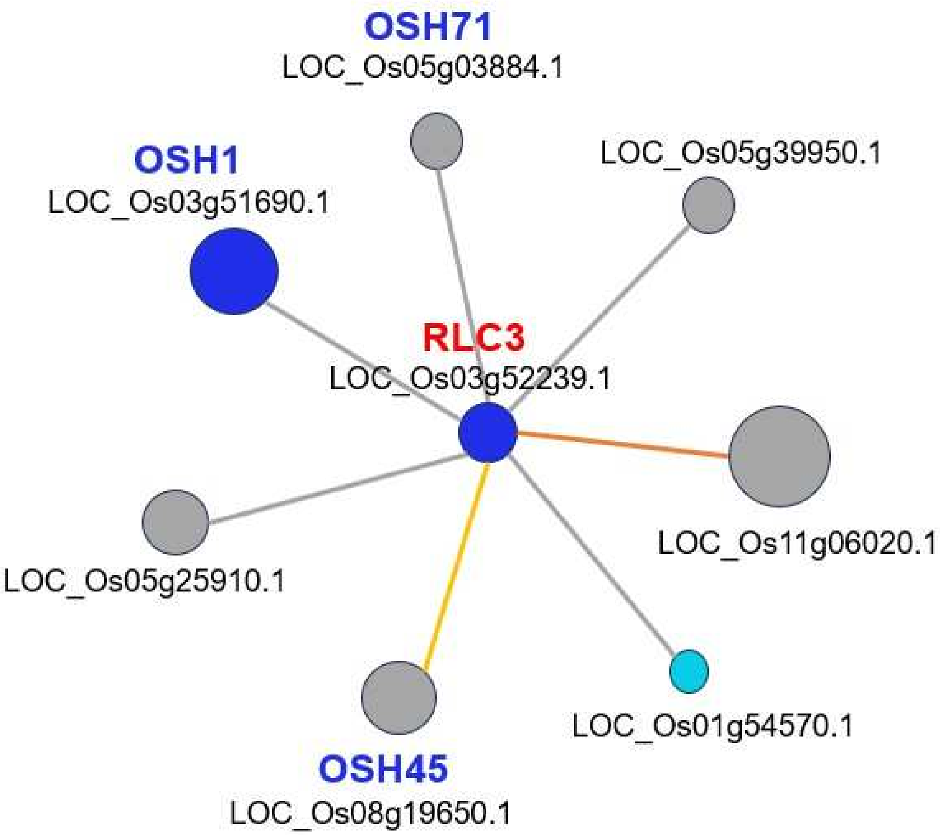


**Fig. S13.** Prediction of RLC3-interacting proteins using the PRIN database.

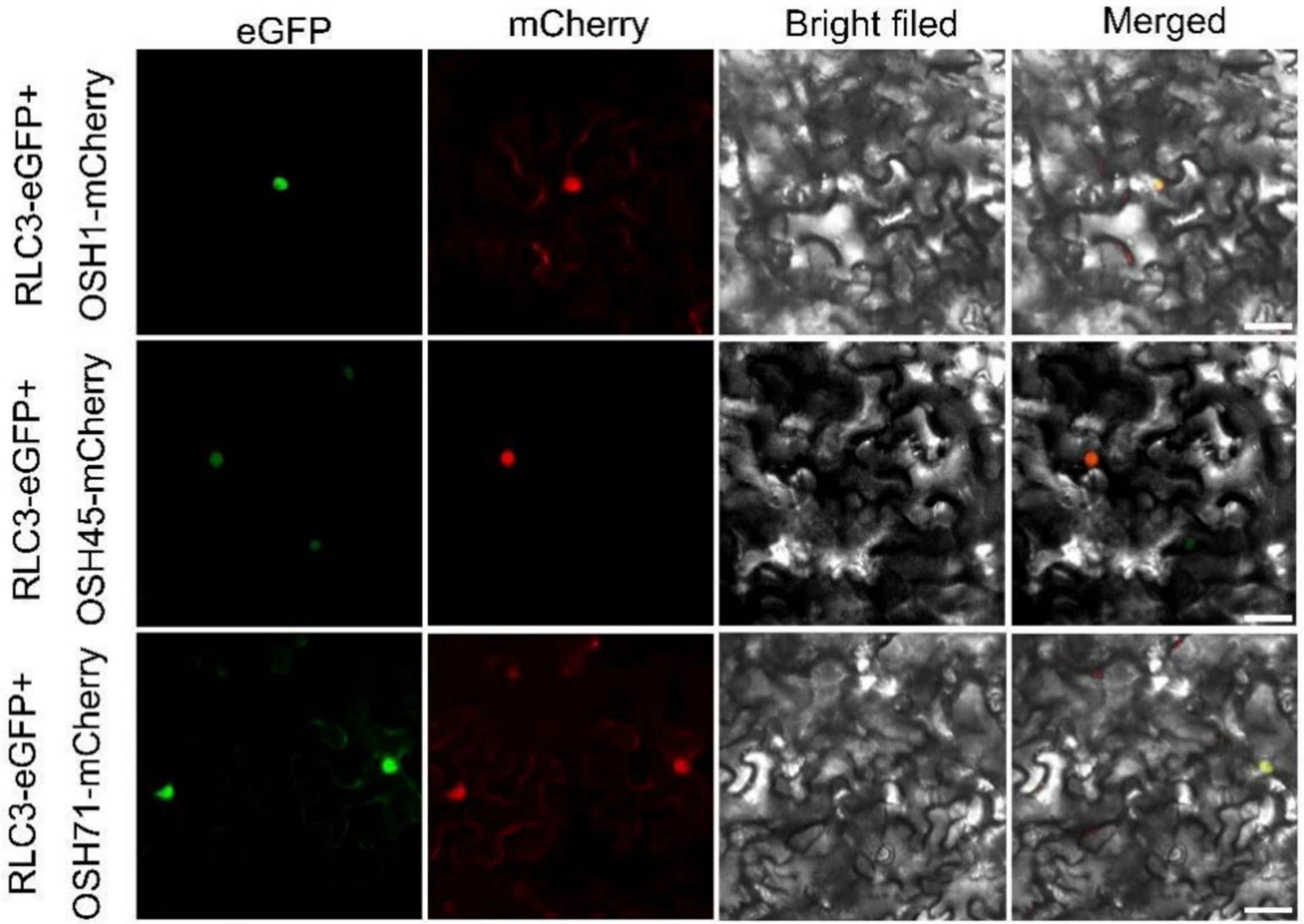


**Fig. S14.** Subcellular co-localization analysis of RLC3 with OSH1, OSH45, and OSH71 proteins. eGFP indicates the green fluorescence channel at 488 nm, mCherry indicates to the red fluorescence channel at 561 nm, Bright field indicates the field of view, Merged refers to the result of three channels overlapping each other. bars=50 μm.

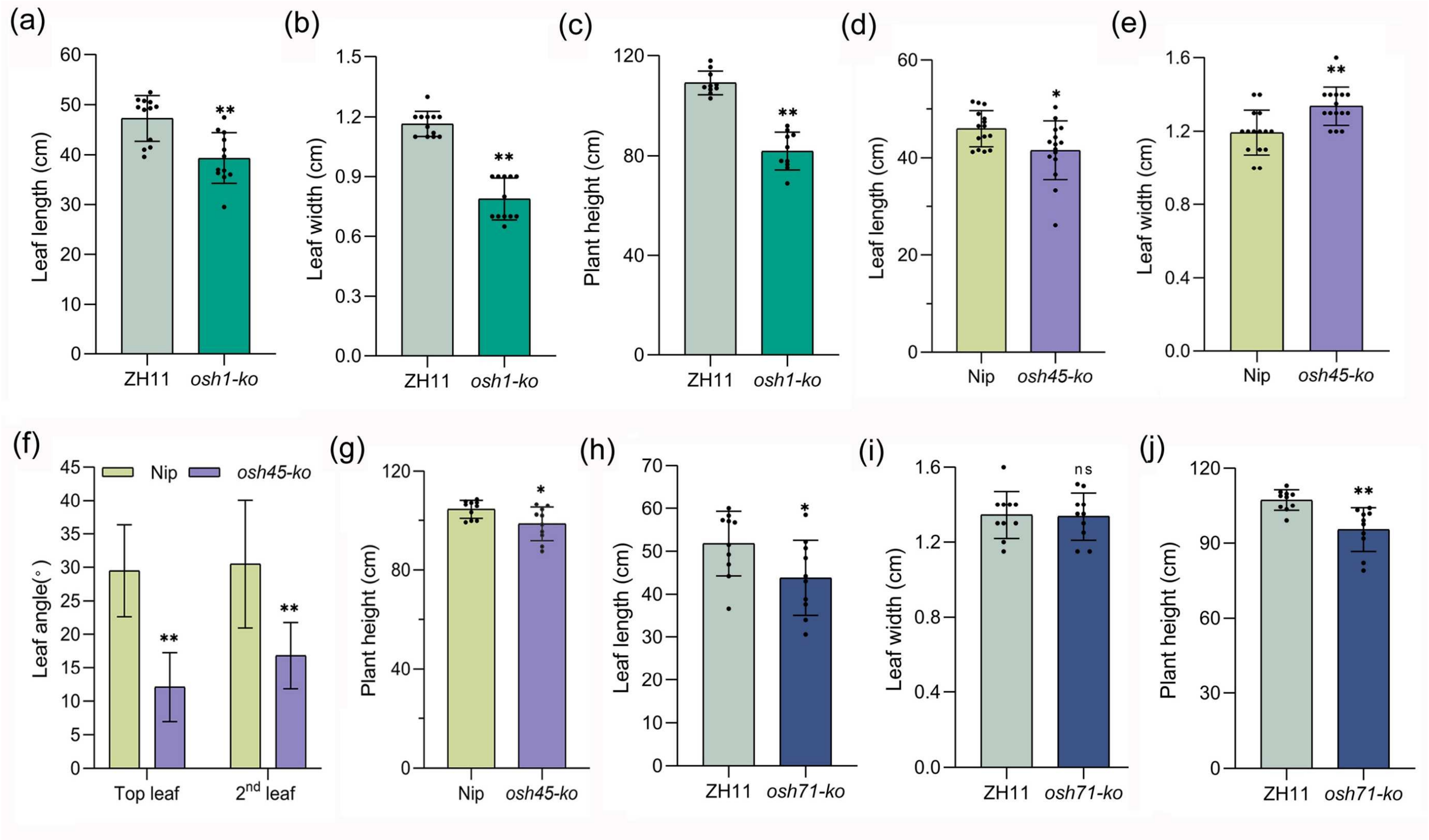


**Fig. S15.** Comparison of flag leaf length, width, and plant height at maturity between wild-type and *OSH1*, *OSH45*, and *OSH71* knockout mutants. (a) Leaf length of wild-type and *OSH1* knockout mutant, n=10. (b) Leaf width of wild-type and *OSH1* knockout mutant, n=10. (c) Plant height of wild-type and *OSH1* knockout mutant, n=10. (d) Leaf length of wild-type and *OSH45* knockout mutant, n=15. (e) Leaf width of wild-type and *OSH45* knockout mutant, n=15. (f) Leaf angle of wild-type and *OSH45* knockout mutant, n=10. (g) Plant height of wild-type and *OSH45* knockout mutant, n=10. (h) Leaf length of wild-type and *OSH71* knockout mutant, n=10. (i) Leaf width of wild-type and *OSH71* knockout mutant, n=10. (j) Plant height of wild-type and *OSH71* knockout mutant, n=10. Data are means ± SD, asterisks indicate significant differences according to Student's *t*-test (*$P < 0.05$; **$P < 0.01$, ns, no significant difference).

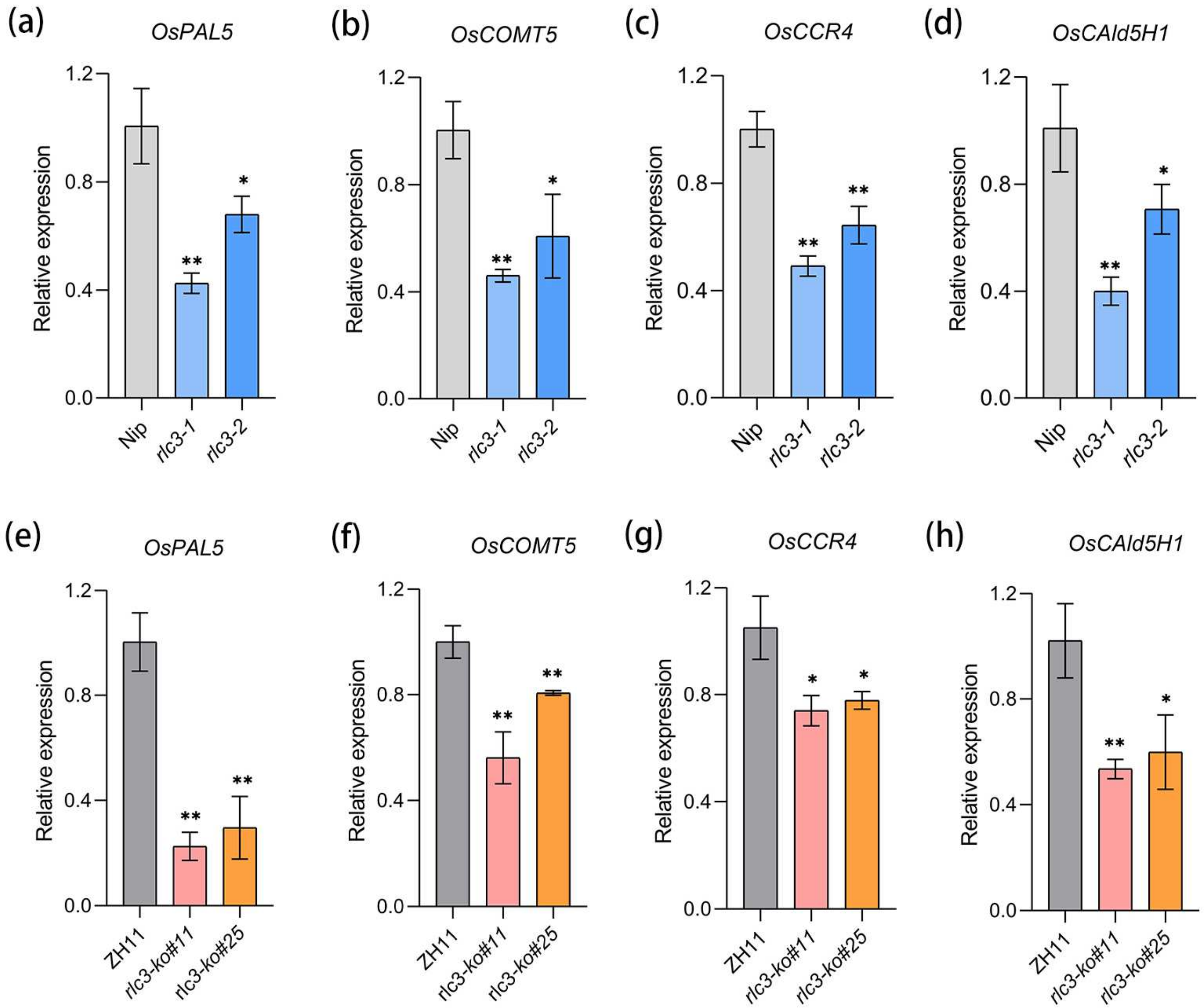


**Fig. S16.** Expression analysis of lignin biosynthesis genes (*OsPAL5*, *OsCOMT5*, *OsCCR4*, and *OsCAld5H1*) in wild-type and *rlc3* mutant lines. (a-d) Expression analysis of lignin biosynthesis genes (*OsPAL5*, *OsCOMT5*, *OsCCR4*, and *OsCAld5H1*) in wild-type Nip, *rlc3-1, and rlc3-2* mutants. (e-h) Expression analysis of lignin biosynthesis genes (*OsPAL5*, *OsCOMT5*, *OsCCR4*, and *OsCAld5H1*) in wild-type ZH11, *rlc3-1, and rlc3-2* mutants. The *Ubiquitin* (*Ubq*) gene is used as a control. Data are means ± SD of three biological replicates; asterisks indicate significant differences according to Student's *t*-test (*$P < 0.05$; **$P < 0.01$).
